# Disruption of the SAGA CORE triggers collateral degradation of KAT2A

**DOI:** 10.1101/2025.07.24.666361

**Authors:** Paul Batty, Hannah Beneder, Caroline Schätz, Gabriel Onea, Maciej Zaczek, Ana P. Kutschat, Sophie Müller, Giulio Superti-Furga, Georg E. Winter, Davide Seruggia

**Affiliations:** St. Anna Children’s Cancer Research Institute (CCRI), Vienna, Austria; CeMM Research Center for Molecular Medicine, Austrian Academy of Sciences, Vienna, Austria; Department of Pediatrics and Adolescent Medicine, Medical University of Vienna, Vienna, Austria; Center for Physiology and Pharmacology, Medical University of Vienna, Vienna, Austria; AITHYRA Institute for Biomedical Artificial Intelligence, Vienna, Austria

**Keywords:** <u>Keywords</u>: SAGA, KAT2A, histone acetyltransferase, collateral degradation, UBR5, proteasome

## Abstract

The SAGA transcriptional co-activator complex regulates gene expression through histone acetylation at promoters, mediated by its histone acetyl transferase, KAT2A. While its structure and function have been extensively investigated, how the stability of individual subunits of SAGA, including KAT2A, is regulated, remains unclear. Here, using a fluorescence-based KAT2A stability reporter, we systematically dissect the molecular dependencies controlling KAT2A protein abundance. We identify the non-enzymatic SAGA CORE module subunits—TADA1, TAF5L, and TAF6L— as necessary for KAT2A stability, with loss of these subunits disrupting the integrity of SAGA, leading to non-chromatin-bound KAT2A that is degraded by the proteasome, consequently leading to reduced H3K9 acetylation. Proteomic profiling reveals progressive loss of CORE and HAT components upon acute disruption of the SAGA CORE, indicating that an intact CORE is required for the stability of numerous components of SAGA. Finally, a focused CRISPR screen of ubiquitin-proteasome system genes identifies the E3 ligase UBR5, a known regulator of orphan protein degradation, and the deubiquitinase OTUD5, as regulators of KAT2A degradation when the SAGA CORE is perturbed. Together, these findings reveal a dependency of KAT2A protein stability on SAGA CORE integrity and define an orphan quality control mechanism targeting unassembled KAT2A, revealing a potential vulnerability in SAGA-driven malignancies.

## INTRODUCTION

Gene expression is controlled by large multi-protein complexes that interact with chromatin and transcription factors to facilitate or repress transcription (Chen & Dent, 2014). The biogenesis and maintenance of such multi-protein complexes is a highly regulated process (Ahnert *et al*, 2015; Natan *et al*, 2017), with ribonucleoprotein immunoprecipitation assays demonstrating that assembly of large protein complexes is often co-translational (Natan *et al*, 2017; Bernardini & Tora, 2024). For instance, components of COMPASS (Halbach *et al*, 2009), TFIID (Bernardini *et al*, 2023), and other co-activator complexes including ATAC and SAGA (Yayli *et al*, 2023) display paired interactions in the cytosol already at the stage of nascent protein, suggesting that components of those complexes are assembled co-translationally. This mechanism is sought to extend the half-life of intrinsically unstable components, with the partner protein acting as a chaperone for aggregation-prone subunits. Numerous quality control mechanisms exist to maintain homeostasis in protein complex formation. These mechanisms are crucial to ensure that proteins fold correctly and are expressed in the right amounts, thereby preventing the formation of mislocalised, misfolded, or aggregated proteins which would have deleterious consequences for the cell (Padovani *et al*, 2022; Pla-Prats & Thomä, 2022). However, as protein complex assembly is not a stoichiometric process and complex partners have different abundancies in the cell, some subunits therefore remain uncomplexed, as there are insufficient binding partners to successfully form the full complex. These ‘orphan’ proteins may become bound by chaperones until such a time that a partner protein becomes available, but in the absence of their normal partners can also misfold. Misfolding can in turn expose hidden degrons or hydrophobic patches, which ordinarily are inaccessible in the assembled complex (Kong *et al*, 2021). These de-novo degrons then mediate protein degradation via specialised machinery of the orphan quality control system, thereby maintaining protein homeostasis (Xu *et al*, 2016; Yanagitani *et al*, 2017; Mark *et al*, 2023).

Interestingly, inherent features of these two mechanisms, i.e. dependency on co-translational assembly and dedicated orphan quality control, can be exploited using chemical or genetic perturbations. For example, chemical (Hsu *et al*, 2020) or genetic (Leeb *et al*, 2010; Xu *et al*, 2015) perturbation of the PRC2 component EED leads to loss of its complex partners, EZH2 and SUZ12, in a phenomenon termed ‘collateral degradation’ (Zhang *et al*, 2023). Similarly, PROTACs targeting HDAC1/2 result in the degradation of other proteins in the same complex (Xiong *et al*, 2021), such as LSD1 of CoREST, or components of the SIN3 complex (Smalley *et al*, 2022), while genetic depletion of HDAC1/2 in neuroblastoma cells leads to destabilisation of NuRD (Zhang *et al*, 2023). Such collateral degradation of partner proteins upon loss of a single complex component has recently emerged as a new therapeutic modality when targeting co-activator or co-repressor complexes that are deregulated in cancer (Mabe *et al*, 2024), as specific targeting of one subunit has the potential to result in collateral loss of multiple subunits, and consequent loss of complex activity.

The SAGA (Spt-Ada-Gcn5 acetyltransferase) is a large co-activator complex composed of 20 proteins organised into five distinct modules (Herbst *et al*, 2021). Two of these five modules, the DUB (Deubiquitinase) and HAT (Histone Acetyl transferase), have enzymatic activity, conferred respectively by the histone deubiquitinase USP22, which deubiquitinates Lysine 120 of Histone H2B, or the acetyltransferase KAT2A (also known as GCN5) (Dent, 2024), which predominately acetylates Histone H3K9 at promoters to facilitate transcriptional activation. The other three modules lack enzymatic activity and have various functions, from mediating interactions with transcription factors (TRRAP), associating with splicing and DNA repair factors (SPL), or serving as a structural scaffold for the other modules (CORE).

Due to their key role in transcriptional activation, acetyltransferases such as KAT2A are desirable therapeutic targets, particularly in cancer (Mustachio *et al*, 2020; White *et al*, 2024). Indeed, KAT2A has been identified as a specific vulnerability in acute myeloid leukemia (AML) (Tzelepis *et al*, 2016), while numerous studies have reported an interplay between KAT2A and MYC-driven oncogenic transcriptional programmes across different cancer subtypes (Mustachio *et al*, 2019; Farria *et al*, 2019, 2020; Malone *et al*, 2024; Liu *et al*, 2025). Multiple independent genetic screens have also identified cancer vulnerabilities within non-enzymatic components of SAGA (Xu *et al*, 2018; Durbin *et al*, 2018; de Almeida *et al*, 2021; Barbosa *et al*, 2024; Chan *et al*, 2024), underscoring their functional importance and showing that targeting these components also holds therapeutic potential.

Notably, inactivation of CORE and HAT components other than KAT2A often phenocopies loss of KAT2A itself (Chan *et al*, 2024; Malone *et al*, 2024), while several studies have reported a reduction in H3K9ac levels upon loss of both HAT and non-HAT SAGA components (Seruggia *et al*, 2019; Fischer *et al*, 2021; Malone *et al*, 2024; Barbosa *et al*, 2024), suggesting that KAT2A’s acetyltransferase activity is dependent on protein-protein interactions within SAGA. However, the contribution of individual SAGA subunits to the regulation of KAT2A protein function and stability still remains unclear.

Here, we used a fluorescence-based stability reporter to systematically measure KAT2A protein levels upon genetic perturbation of SAGA components. We found that specific components of the structural CORE module of SAGA are necessary for KAT2A protein stability, recruitment to promoters and acetylation of histones. Fractionation experiments revealed loss of complex integrity and disengagement of the HAT module upon SAGA CORE disruption, resulting in the accumulation of non-complexed KAT2A protein that is prone to proteasome-mediated degradation. Using pooled CRISPR screening we identified proteins that regulate KAT2A turnover upon perturbation of the SAGA CORE, including OTUD5 and UBR5, components of the orphan quality control system, implicating them as effectors of solitary KAT2A protein degradation.

## RESULTS

### Arrayed CRISPR screening to monitor KAT2A stability upon knockout of SAGA components

To monitor KAT2A protein abundance upon genetic or chemical perturbations, we established a stability reporter by fusing the *KAT2A* coding sequence in-frame with BFP, followed by mCherry, with the two fluorophores separated by a P2A peptide linker. In this system BFP therefore reports on KAT2A abundance while mCherry serves as a normalisation control, in a setup amenable to pooled or arrayed CRISPR screening using flow cytometry (de Almeida *et al*, 2021; Schwartz *et al*, 2023; Hsia *et al*, 2024). We expressed the KAT2A stability reporter in wild type HAP1 cells along with Cas9 and performed arrayed CRISPR screening using guide RNAs (gRNAs) targeting the 20 components of the SAGA complex (**Fig. 1A, B**), along with positive and negative controls **(Table S1).** Each gRNA also expressed GFP, allowing discrimination of transduced from non-transduced cells in a heterogenous cell population, in addition to guide RNA dropout over time by comparing the percentage of GFP positive cells for each guide at each time point of the experiment. As expected, due to their common essentiality, the GFP signal originating from gRNAs targeting the essential genes *RAD21* and *MYC* was rapidly lost, in addition to the *SF3B3* and *SF3B5* components of the SAGA splicing module (SPL) (**Fig. S1A-C**). Thus, wild type HAP1^Cas9^ have high levels of Cas9 activity and allow efficient knockout of target genes and are therefore amenable to arrayed CRISPR screening.

**Figure 1.**
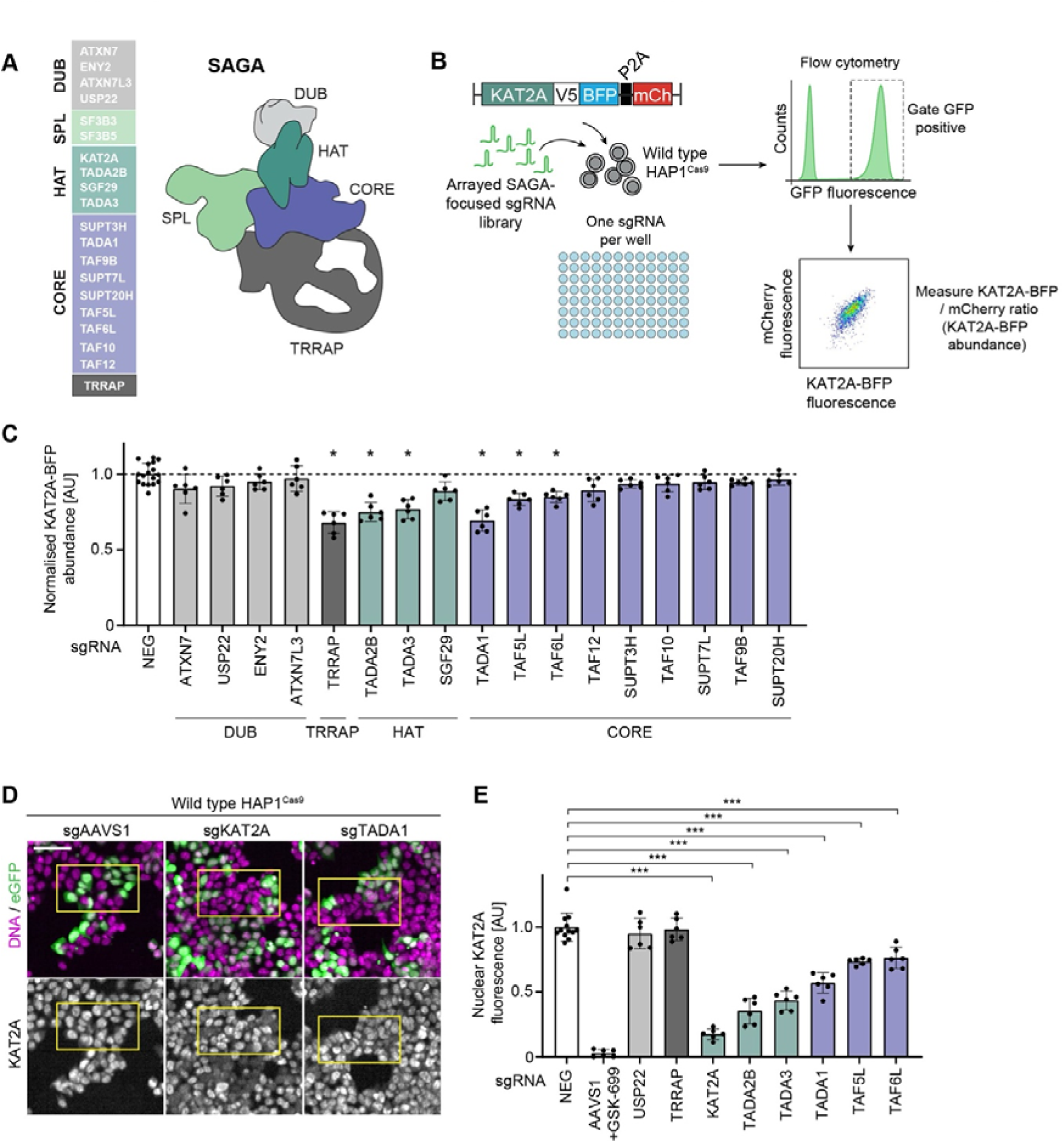
Arrayed CRISPR screening to monitor KAT2A stability upon knockout of SAGA components. **A)** Structure of the SAGA complex. The SAGA complex is composed of 20 proteins arranged into five distinct modules. **B)** Schematic of the KAT2A stability reporter and workflow of the arrayed CRISPR screen. BFP was fused in frame with the KAT2A open reading frame in the stability reporter, followed by a P2A sequence and mCherry tag for normalisation. Wild type HAP1^Cas9^ cells expressing the KAT2A stability reporter were transduced with an arrayed SAGA-focused gRNA library, with one gRNA transduced per well. GFP positive cells were gated and the BFP/mCherry ratio measured for each guide in the library. **C)** Arrayed CRISPR screen of KAT2A-BFP protein abundance following knockout of SAGA subunits. Wild type HAP1^Cas9^ cells expressing the KAT2A stability reporter were transduced with GFP-expressing gRNAs targeting each of the 20 components of SAGA, as well as control gRNAs. 5 days post-transduction, the KAT2A-BFP levels of GFP positive cells was quantified using flow cytometry. Dots represent the mean normalised KAT2A-BFP fluorescence of technical replicates, error bars represent the standard deviation, bars indicate the mean for each condition. One gRNA per SAGA subunit is plotted. Significance was tested using a two-tailed Mann–Whitney U-test. **D-E)** Validation of reduced KAT2A protein abundance following knockout of SAGA CORE and HAT components using high throughput immunofluorescence. **D)** Representative immunofluorescence images of wild type HAP1^Cas9^ cells transduced with GFP expressing gRNAs against *AAVS1*, *KAT2A*, or *TADA1*, as indicated. 5 days post-transduction, cells were fixed and KAT2A was stained with an anti-KAT2A antibody. DNA was stained with Hoechst 33342. **E)** Quantification of nuclear KAT2A fluorescence by immunofluorescence for GFP positive cells for the indicated gRNAs. Dots represent the median nuclear KAT2A fluorescence of technical replicates, error bars indicate the standard deviation, bars indicate the mean for each condition. Significance was tested using a two-tailed Mann–Whitney U-test. Data information: **C)** (*) P < 0.05; two-tailed Mann–Whitney U-test for each of the two independent biological replicates. **E)** (***) P < 0.001; two-tailed Mann–Whitney U-test. Biologically independent replicates: **(C-E)** (n = 2). In **C, E,** only GFP positive cells were analysed, and data was normalised relative to sgAAVS1 transduced cells treated with the KAT2A/KAT2AB PROTAC GSK-699 (100 nM) for 6 h. All images show single Z-slices. Yellow boxes indicate inset regions showing non-transduced (GFP negative) and transduced (GFP positive) cells. Scale bars: **D,** 50 µm. The number of cells analysed by microscopy in **E** is listed in **Table S3.**

To validate that the KAT2A stability reporter was responsive to changes in KAT2A protein abundance, we treated HAP1^Cas9^ cells with GSK-699, a potent, recently developed KAT2A/KAT2B PROTAC (Bassi *et al*, 2018; Malone *et al*, 2024). Following 6 h treatment with the PROTAC, we observed a strong reduction in KAT2A-BFP signal but no change in mCherry fluorescence (**Fig. S1D-F**), as expected as mCherry is translated as a separate polypeptide, with a > 80 % reduction in KAT2A-BFP fluorescence for cells treated with the PROTAC compared to DMSO-treated controls (from 1.0 ± 0.07 (DMSO) to 0.19 ± 0.09 (GSK-699), mean + S.D.). Additionally, cells transduced with gRNAs targeting *KAT2A* itself had substantially reduced KAT2A-BFP and mCherry mean fluorescence, with a > 50 % reduction in signal for both fluorophores compared to *AAVS1* transduced controls **(Fig. S1G, H)**. Thus, the KAT2A stability reporter construct faithfully reports on KAT2A protein levels upon chemical or genetic perturbation. To systematically investigate the role of SAGA components on KAT2A abundance, we gated GFP positive cells and measured the KAT2A-BFP/mCherry ratio for each gRNA in our collection (**Fig. 1C)** relative to negative control guides targeting either the *AAVS1* safe harbour locus or non-targeting sequences. Targeting *ATXN7*, *ATXN7L3*, *USP22* and *ENY2*, components of the DUB module, did not significantly alter KAT2A levels, consistent with the observation that the DUB module is loosely bound to SAGA and hence dispensable for its structural integrity (Herbst *et al*, 2021). However, knockout of *TADA2B* and *TADA3*, components of the HAT module that are in close proximity to KAT2A within SAGA, resulted in a significant drop in KAT2A abundance, indicating that KAT2A protein levels are reduced upon loss of its partners in the HAT module. Targeting *TRRAP*, the largest subunit of SAGA, also resulted in reduced KAT2A abundance using the stability reporter assay. While not structurally close to the HAT module (Herbst *et al*, 2021), it has been reported that TRRAP and KAT2A engage in a tertiary structure with MYC (McMahon *et al*, 2000; Liu *et al*, 2003) which might become disrupted upon loss of *TRRAP*. However, closer inspection of the 2D FACS plots for cells transduced with guides against *TRRAP* showed that these cells had an increase in mCherry signal compared to control cells, rather than a decrease in KAT2A-BFP fluorescence (**Fig. S1I**), suggesting an artefact of the reporter leads to a reduced KAT2A-BFP/mCherry ratio when targeting *TRRAP.* Interestingly, upon targeting the non-enzymatic CORE module of SAGA, we identified three subunits, *TADA1*, *TAF5L* and *TAF6L*, whose knockout also resulted in significantly reduced levels of KAT2A (**Fig. 1C**). These results therefore suggest that selected CORE components of SAGA, in addition to components of the HAT module, play a role in regulating KAT2A protein abundance.

To validate these findings, we turned to high-throughput confocal microscopy and measured endogenous KAT2A levels by immunofluorescence in wild type HAP1^Cas9^ cells, following knockout of SAGA subunits that reduced KAT2A abundance in the stability reporter screen. Guides against *KAT2A* itself or treatment with GSK-699 served as positive controls, with guides against *AAVS1* used as a negative control. As in the arrayed screen, each gRNA also expressed a GFP cassette. Thus, by thresholding the nuclei of GFP positive cells, we could specifically measure the nuclear KAT2A fluorescence of transduced cells for each guide (**Fig. 1D, E**). Importantly, we observed that knockout of KAT2A yielded a drop in KAT2A fluorescence comparable to that induced by the PROTAC, demonstrating the sensitivity of the assay (**Fig. 1D, E**). Knockout of *TRRAP* had no effect on endogenous KAT2A abundance, confirming that its effect on the stability reporter in the arrayed screen was an artefact of the reporter, while loss of *USP22* also did not reduce KAT2A levels. Consistent with our screen findings, knockout of the HAT subunits *TADA2B* and *TADA3*, as well as the CORE subunits *TADA1*, *TAF5L* and *TAF6L* resulted in a significant reduction in nuclear KAT2A protein levels (**Fig. 1D, E)**, confirming that components of the HAT module, in addition to non-enzymatic structural components of SAGA regulate KAT2A protein abundance. Importantly, targeting of other CORE subunits such as *SUPT20H*, *SUPT3H*, or *TAF10*, did not significantly reduce KAT2A protein levels (**Fig. S1J**), thereby demonstrating a specific effect upon targeting *TADA1, TAF5L,* and *TAF6L.* Altogether, these results demonstrate that KAT2A protein levels are reduced upon loss of its binding partners in the HAT module, but also upon knockout of selected components of the non-enzymatic CORE, which do not interact with KAT2A directly but instead play a structural role in the complex.

### Components of the SAGA CORE regulate KAT2A abundance and HAT function

To mechanistically dissect KAT2A loss upon targeting the SAGA CORE, we focused on the three CORE components identified in the arrayed CRISPR screen and generated homozygous HAP1 knockout cell lines for *TAF5L*, *TAF6L* and *TADA1*, in addition to *KAT2A* to serve as a positive control. Clonal TAF5L, TAF6L and TADA1 KO cells showed a drastic reduction in endogenous nuclear KAT2A fluorescence, with a reduction in KAT2A fluorescence comparable to that of KAT2A KO cells (**Fig. S2A, B**), with a similar reduction in nuclear KAT2A protein levels observed across multiple independent knockout clones (**Fig. S2B**), confirming our previous findings with knockout pools. Importantly, overexpression of murine full length *Tada1, Taf5l* or *Taf6l* cDNA in the corresponding KO clone restored *KAT2A* fluorescence to wild type levels (**Fig. 2A, B**). Thus, TAF5L, TAF6L, TADA1 are necessary to maintain *KAT2A* protein levels.

**Figure 2.**
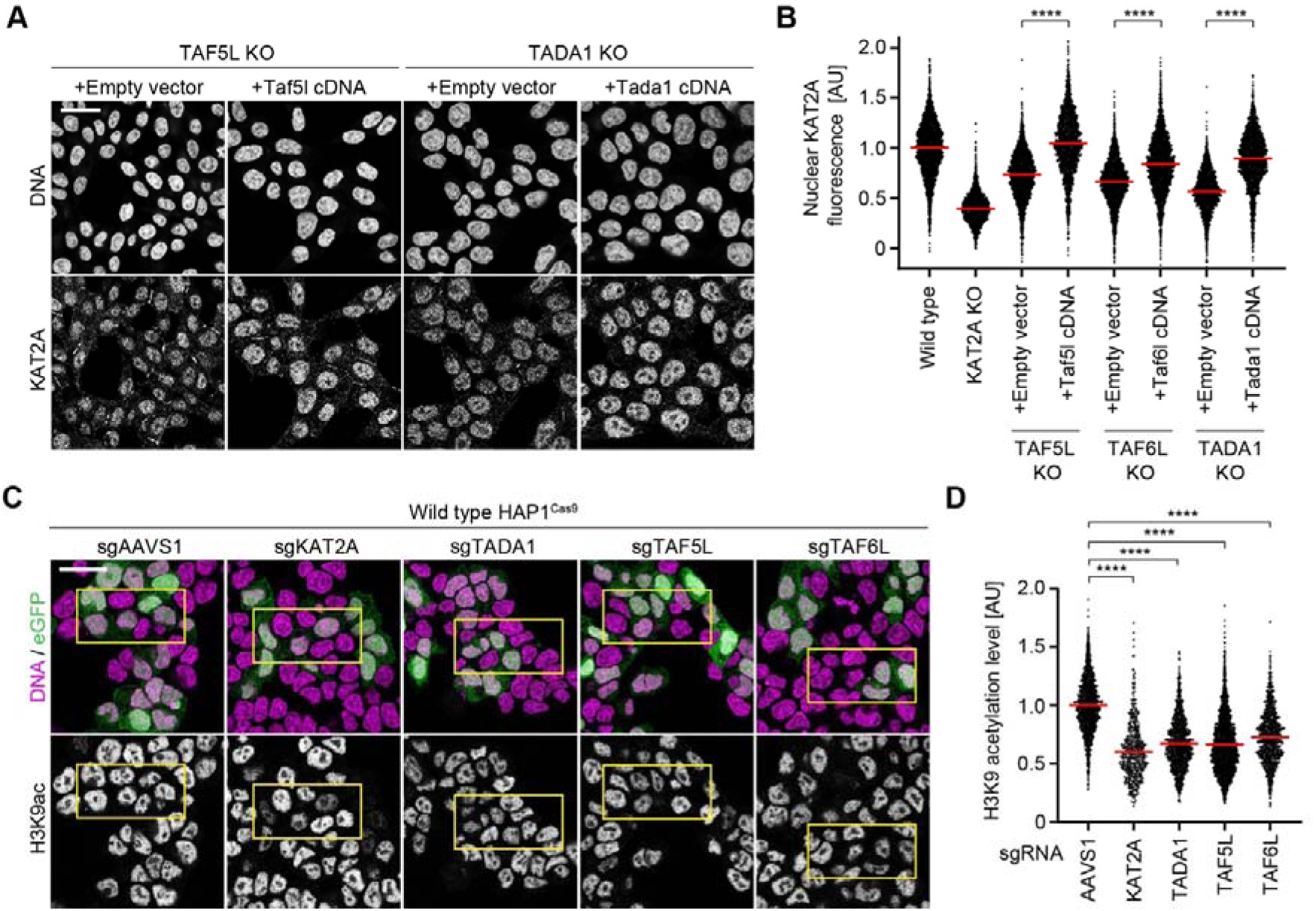
Components of the SAGA CORE regulate KAT2A protein abundance and HAT function. **A, B)** Overexpression of full-length cDNA for *TAF5L*, *TAF6L*, or *TADA1* restores nuclear KAT2A fluorescence to wild type levels in the respective knockout cell line. **A)** Representative immunofluorescence images of TAF5L KO or TADA1 KO HAP1 cells, stably overexpressing either an empty vector plasmid or full-length murine *Taf5l* or *Tada1* cDNA. Cells were fixed for immunofluorescence and KAT2A was stained with an anti-KAT2A antibody. DNA was stained with Hoechst 33342. **(B)** Quantification of nuclear KAT2A fluorescence by immunofluorescence for TAF5L KO, TAF6L KO, and TADA1 KO cells stably overexpressing either an empty vector plasmid or full-length murine *Taf5l*, *Taf6l*, or *Tada1* cDNA for the respective cell line. Dots represent the mean nuclear KAT2A fluorescence of individual cells; red bars indicate the median. Significance was tested using a two-tailed Mann–Whitney U-test. **C-D)** Global H3K9 acetylation levels are strongly reduced following knockout of the SAGA CORE components TADA1, TAF5L, or TAF6L. **C)** Representative immunofluorescence images of wild type HAP1^Cas9^ cells transduced with indicated GFP expressing gRNAs. 6 days post-transduction, cells were fixed and H3K9ac was stained with an anti-H3K9ac antibody. DNA was stained with Hoechst 33342. **D)** Quantification of H3K9 acetylation signal by immunofluorescence for GFP positive cells. The H3K9ac acetylation level was calculated by dividing the mean H3K9ac fluorescence for each cell by the DNA fluorescence as measured by Hoechst 33342 staining. Dots represent the mean H3K9ac acetylation level of individual cells; red bars indicate the mean. Significance was tested using a two-tailed Mann–Whitney U-test. Data information: **B, D** (****) P < 0.0001; two-tailed Mann–Whitney U-test. Biologically independent replicates: **A-B** (TAF5L KO (n = 3), TAF6L KO (n = 6), wild type (n = 7), KAT2A KO (n = 7), TADA1 KO (n = 4). **C, D** (n = 3). In **B** the data was normalised relative to the median nuclear KAT2A fluorescence of wild type HAP1 cells. In **D** only GFP positive cells were analysed. All images show single Z-slices. Yellow boxes indicate inset regions showing non-transduced (GFP negative) and transduced (GFP positive) cells. Scale bars: 20 µm. The number of cells analysed by microscopy in **B, D** is listed in **Table S3.**

To address which specific domains within the SAGA CORE play a role in regulating KAT2A protein abundance, we turned to publicly available CryoEM data (Herbst *et al*, 2021) to try and identify candidate domains that might mediate protein-protein interactions within SAGA that regulate complex integrity, which might be necessary for maintaining KAT2A protein levels. Based on their position in the complex bridging the HAT and CORE modules, we hypothesised that the six WD40 repeats of TAF5L, the HEAT repeat of TAF6L as well as an intrinsically disordered region (IDR) at the C-terminus of the protein might be relevant for their function. Indeed, overexpression of cDNA constructs lacking TAF5L WD40 domains, TAF6L HEAT repeats, or the C-terminal IDR of TAF6L failed to rescue KAT2A levels, although a construct lacking the N-terminal domain of TAF5L rescued nuclear KAT2A fluorescence to the same extent as the full-length construct, and is therefore dispensable for regulating KAT2A protein abundance **(Fig. S2C-H).** Thus, specific domains involved in protein-protein interactions of SAGA CORE components are both necessary and sufficient to maintain KAT2A protein levels.

Having established that loss of *TAF5L, TAF6L,* or *TADA1* reduces *KAT2A* protein levels, we wished to address if their loss also resulted in global changes in H3K9 acetylation, the main histone mark deposited by KAT2A on chromatin. To quantitatively address this question, we turned to confocal microscopy and measured H3K9ac fluorescence relative to a DNA counterstain in HAP1 knockout pools following gRNA transduction, and thresholded GFP positive nuclei to identify transduced cells. Indeed, targeting *TADA1*, *TAF5L* or *TAF6L* led to a strong reduction in global H3K9ac levels compared to *AAVS1-*transduced controls (**Fig. 2C, D**), consistent with previous findings in mouse embryonic stem cells (Seruggia *et al*, 2019). Interestingly, the three SAGA CORE mutants displayed a global reduction in H3K9 acetylation comparable to that of KAT2A knockout cells (relative H3K9ac signal: *sgKAT2A:* 0.60 ± 0.24, *sgTAF5L:* 0.66 ± 0.24, *sgTAF6L:* 0.73 ± 0.23, *sgTADA1:* 0.67 ± 0.22, mean + S.D.). Thus, despite residual amounts of KAT2A protein in the nucleus in CORE KO cells (**Fig. 2B, S2B**), KAT2A’s histone acetylation output is substantially reduced upon loss of *TAF5L, TAF6L, or TADA1*, with H3K9 acetylation reduced to the same extent as in cells lacking KAT2A when the SAGA CORE is perturbed.

### Loss of TAF5L and TADA1 leads to HAT module disengagement from SAGA

As perturbing the SAGA CORE phenocopies loss of KAT2A at the level of global H3K9 acetylation, we hypothesised that recruitment of KAT2A to chromatin is abrogated in SAGA CORE mutants. To test this hypothesis, we therefore measured KAT2A chromatin occupancy by CUT&RUN (Skene & Henikoff, 2017). While we could detect enrichment of KAT2A signal around transcription start sites in wild type cells, knockout of *TAF5L* phenocopied loss of *KAT2A*, as we were unable to detect KAT2A on chromatin in neither KAT2A nor TAF5L KO cells (**Fig. 3A, B**). Thus, not only is KAT2A protein abundancy reduced upon perturbing the SAGA CORE, but the residual protein cannot be recruited to its canonical binding sites on chromatin around promoters.

**Figure 3.**
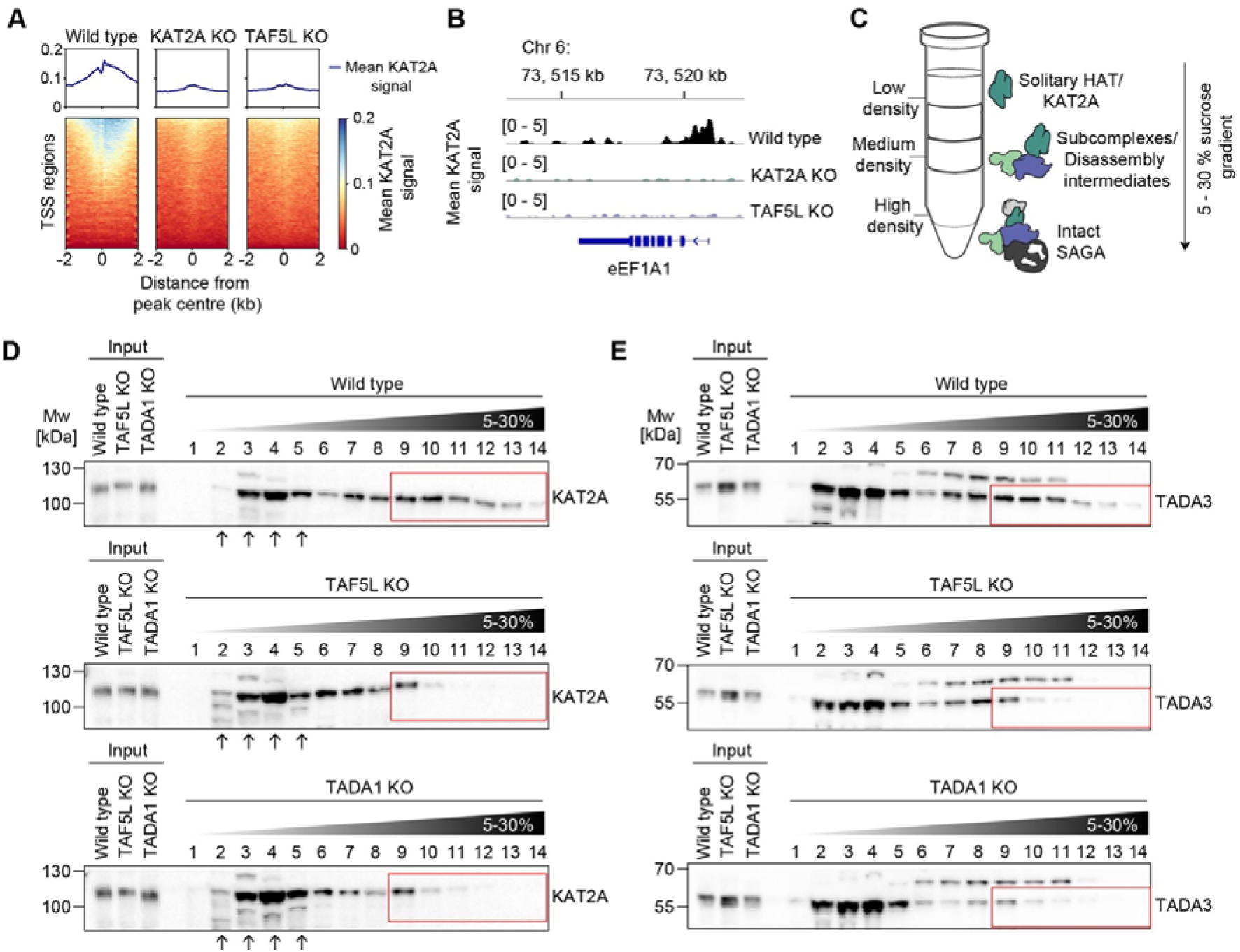
Loss of TAF5L and TADA1 promotes HAT module disengagement from SAGA. **A)** Loss of TAF5L phenocopies loss of KAT2A at the chromatin level. The average KAT2A signal around transcriptional start sites (TSS) was determined by CUT&RUN. Mean KAT2A signal was piled up in a 4 kb window centred around TSS. Line profiles indicate the mean KAT2A signal within the window of interest. **B)** Genomic tracks of KAT2A signal for wild type, KAT2A KO, and TAF5L KO HAP1 cells. The eEF1A1 locus is shown as a representative locus. **C)** Schematic of expected distribution of SAGA complex components following sucrose gradient ultracentrifugation. High-molecular weight species sediment into the more dense sucrose. Subcomplexes or disassembly intermediates of medium density can be found in the middle fractions, while low molecular weight species remain near the top of the gradient. **D-E)** Sucrose gradient fractionation reveals that the SAGA histone acetyl transferase (HAT) disengages from SAGA when the CORE is perturbed. Cell lysates were prepared from wild type, TAF5L KO, and TADA1 KO cells, and loaded onto 5 – 30 % sucrose gradients. Gradients were ultracentrifuged and fractionated to separate protein complexes based on their sedimentation. The resultant fractions were loaded onto an SDS-PAGE gel and analysed by Western blotting, using antibodies against KAT2A **(D)**, or TADA3 **(E**) as indicated. Membranes were developed simultaneously to allow cross-comparison between genotypes. Inputs indicate the cell lysates for each condition prior to ultracentrifugation. Numbers indicate each fraction, moving from lowest to highest density sucrose fractions (5 – 30 %). Red boxes indicate fractions containing high-molecular weight KAT2A or TADA3 species that are retained in wild type cells and absent in TAF5L KO or TADA1 KO cells. Arrows indicate fractions containing low-molecular weight KAT2A species, which are enriched when the SAGA CORE is perturbed. **D)** Immunoblot analysis of KAT2A following sucrose gradient fractionation for wild type, TAF5L KO, and TADA1 KO cells, as indicated. **E)** Immunoblot analysis of TADA3 for wild type, TAF5L KO, and TADA1 KO cells following sucrose gradient fractionation, as in **D**. Data information: Biologically independent replicates: **A, B, D, E** (n = 2).

SAGA CORE perturbation has previously been reported to lead to impaired SAGA complex assembly (Fischer *et al*, 2021; Hisler *et al*, 2024). However, how the HAT module interacts with the CORE and if an intact CORE is required for engagement of the HAT with SAGA is not well characterised. As we did not detect chromatin-bound KAT2A at promoters upon targeting the SAGA CORE, we therefore reasoned that TAF5L, TAF6L, and TADA1 might mediate important protein-protein interactions within the CORE that are necessary for engagement of the HAT module with the rest of SAGA. In such a situation, loss of these proteins would lead to separation of the HAT module and an increase in non-complexed KAT2A protein that can no longer bind to chromatin. To test this hypothesis, we turned to sucrose gradient fractionation (**Fig. 3C)**. High-molecular weight species, such as the intact SAGA complex, containing all or most of its components, have high density and thus migrate into the dense sucrose fractions. Disassembly intermediates or subcomplexes of intermediate molecular weight migrate into middle fractions, while solitary proteins or individual complex modules of low molecular weight remain near the top of the gradient **(Fig. 3C).**

To investigate to what extent perturbation of the SAGA CORE affects engagement of the HAT module with SAGA, we therefore ultracentrifuged and fractionated sucrose gradients from wild type, TAF5L KO, and TADA1 KO cells, before western blotting against KAT2A, as well as TADA3, another HAT component. In wild type cells, KAT2A and TADA3 could be detected across the full range of fractions **(Fig. 3D, E),** indicating the presence of both complexed and non-complexed forms of the proteins. Indeed, in wild type cells both KAT2A and TADA3 were readily detectable in high molecular weight fractions (‘heavy fractions’) **(Fig. 3D, E**, red boxes), consistent with their incorporation into fully assembled SAGA complexes, where the HAT module is engaged with the rest of the complex. Strikingly, in the absence of *TAF5L* and *TADA1* we observed a pronounced loss of KAT2A and TADA3 signal in heavy fractions (**Fig. 3D, E**, red boxes), consistent with loss of complex integrity and disengagement of the HAT from the rest of SAGA. We could also detect a concomitant accumulation of KAT2A in lighter fractions in CORE KO cells compared to wild type (**Fig. 3D**, arrows), with this enrichment suggesting a redistribution of KAT2A into low molecular weight species when the CORE is disrupted, either as free-floating protein or as a subcomplex with other HAT components. Thus, together these results demonstrate that TAF5L and TADA1 are not only required to maintain KAT2A protein levels, but are also important regulators of SAGA complex integrity that are necessary to maintain engagement of the HAT module with SAGA.

### Progressive destabilisation of SAGA CORE and HAT upon depletion of TADA1

Constitutive knockout of *TAF5L, TAF6L, TADA1* leads to reduced KAT2A protein abundancy, but as the rate of protein rundown following gene knockout cannot be temporally controlled, in such a setup the kinetics of protein loss cannot be addressed. Therefore, to assess the kinetics of KAT2A protein depletion upon perturbing the SAGA CORE, we expressed a TADA1-FKBP12^F36V^ (dTAG) degron construct (Nabet *et al*, 2018) in TADA1 KO HAP1 cells, using a haemagglutinin (HA) tag used to monitor TADA1 protein levels, with overexpression of *Tada1* restoring nuclear KAT2A levels to that of wild type (**Fig. 2B)**. Immunostaining and confocal microscopy revealed that although TADA1 protein was rapidly degraded within 1 h of dTAG^V^-1 addition (**Fig. 4A, B)**, KAT2A protein levels gradually reduced over time, reaching the level of TADA1 KO cells within 24 h of dTAG^V^-1 treatment (**Fig. 4C, D)**. Thus, acute degradation of TADA1 confirms that KAT2A protein abundance is reduced when the SAGA CORE is perturbed, but that the effect of TADA1 loss on KAT2A protein is not immediate, and that KAT2A instead becomes progressively less abundant over time.

**Figure 4.**
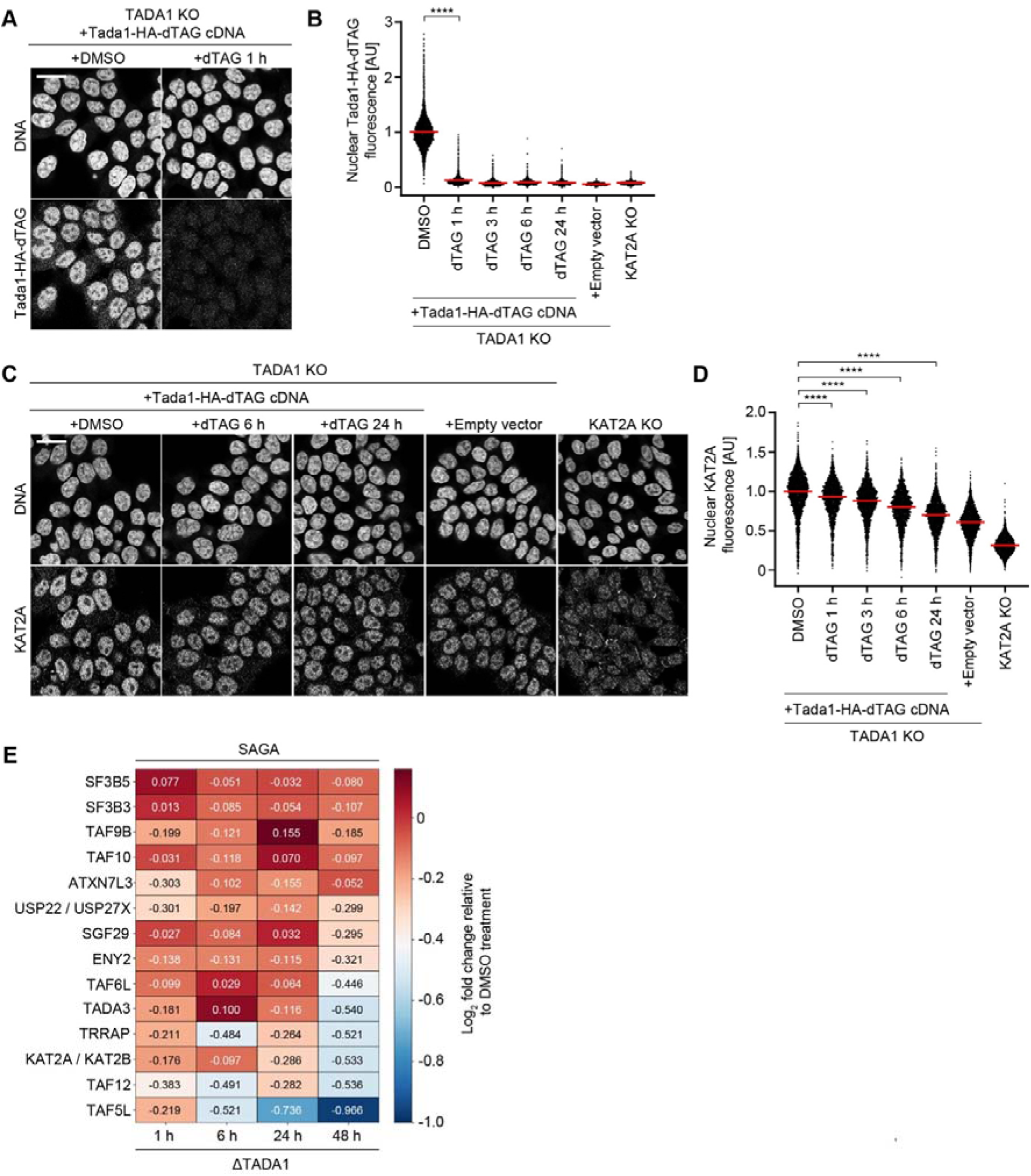
Progressive destabilisation of SAGA CORE and HAT upon depletion of TADA1. **A**, **B**) TADA1-HA-dTAG is efficiently degraded within 1 h. TADA1 KO HAP1 cells overexpressing full length TADA1-HA-dTAG cDNA were treated with DMSO or dTAG^V^-1 as indicated. Following compound treatment, cells were fixed and TADA1-HA-dTAG was stained with an anti-HA antibody. DNA was stained with Hoechst 33342. **B)** Quantification of nuclear TADA1-HA-dTAG fluorescence by immunofluorescence for cell lines in **A)** treated with compounds as indicated. Dots represent the mean nuclear HA fluorescence of individual cells; red bars indicate the mean. Significance was tested using a two-tailed Mann–Whitney U-test. **C, D)** Acute degradation of TADA1 leads to reduced KAT2A abundance. TADA1 KO cells stably overexpressing full-length murine *Tada1* cDNA with a C-terminal FKBP12^F36V^ (dTAG) cassette were treated with DMSO or 100 nM dTAG^V^-1 for the time points as indicated, before fixation for immunofluorescence. **C)** Representative immunofluorescence images of KAT2A KO cells, or TADA1 KO cells overexpressing full length TADA1-dTAG cDNA or an empty vector, treated with DMSO or dTAG^V^-1 as indicated. KAT2A KO and TADA1 KO^empty^ ^vector^ cells were treated with DMSO. Following compound treatment, cells were fixed and KAT2A was stained with an anti-KAT2A antibody. DNA was stained with Hoechst 33342. **D)** Quantification of nuclear KAT2A fluorescence by immunofluorescence for cell lines in **C)** treated with compounds as indicated. Dots represent the mean nuclear KAT2A fluorescence of individual cells; red bars indicate the median. Significance was tested using a two-tailed Mann–Whitney U-test. **E)** Heatmap of differential expression of SAGA complex components following acute depletion of TADA1 for the indicated time points, as determined by TMT-expression proteomics. Log_2_ fold changes of SAGA subunits relative to DMSO-treated controls are shown for each subunit at each time point. Data information: **B, D** (****) P < 0.0001; two-tailed Mann–Whitney U-test. Biologically independent replicates: **A-E** (n = 3). In **B** the data was normalised relative to the mean nuclear HA fluorescence of DMSO-treated cells. In **D** the data was normalised relative to the median nuclear KAT2A fluorescence of DMSO treated TADA1 KO cells overexpressing full-length Tada1-dTAG cDNA. All images show single Z-slices. Scale bars: 20 µm. The number of cells analysed by microscopy in **B, D** is listed in **Table S3.**

Having observed that KAT2A protein levels are reduced following acute degradation of TADA1, we wished to address the consequences of SAGA CORE perturbation on other components of SAGA, as well as the effect globally on other protein complexes. We therefore used TMT-labelling to perform expression proteomics for 5 time points of interest in triplicate, following TADA1 degradation for either 1, 6, 24, or 48 h, and compared protein abundancies relative to control cells treated with DMSO. More than 7200 proteins were successfully identified in our dataset, including 15 of the 20 components of SAGA (**Fig. 4E, Fig. S3A).** Consistent with our immunofluorescence data **(Fig. 4A, B),** TADA1 was significantly depleted after 1 h of dTAG^V^-1 treatment and remained the most strongly depleted protein at all time points (**Fig. S3B-E).** In the absence of unique KAT2A peptides, we annotated common KAT2A/KAT2B peptides to estimate KAT2A abundance and observed progressive depletion over time (**Fig. 4E**), with the most prominent depletion after 48 h dTAG^V^-1 treatment. In addition, the abundancies of several other SAGA components also progressively decreased over time, most prominently the CORE components TAF5L and TAF12, as well as TRRAP, with TAF6L and the HAT component TADA3 also significantly depleted after 48 h dTAG^V^-1 treatment (**Fig. 4E**, **Fig. S3D, E**). To verify the specificity of this effect, we assessed the abundance of components from other protein complexes that share subunits with SAGA, finding minimal changes in the levels of ATAC and TFIID subunits (**Fig. S3B-E).** Thus, the reduction in protein abundancies of SAGA components is driven specifically by the acute depletion of TADA1.

Interestingly, several of the downregulated components are known to directly interact with or are in close structural proximity to TADA1 within SAGA. Indeed, TAF12 interacts with TADA1 via its histone fold domain (Xu *et al*, 2018), in a process that occurs co-translationally (Yayli *et al*, 2023). Similarly, TRRAP has been suggested to form interactions with TADA1 via a cleft bridging TRRAP and the CORE (Herbst *et al*, 2021), with TAF5L and TAF6L also in close proximity to TADA1 (Herbst *et al*, 2021). As cotranslational assembly is a prominent mechanism by which large multi-protein complexes assemble (Duncan & Mata, 2011; Natan *et al*, 2017; Bernardini & Tora, 2024), loss of direct interactors or neighbouring proteins can lead to destabilisation of other complex members (Zhang *et al*, 2023). Consistent with this finding, we observed loss of both TAF12 and TAF5L at early time points after TADA1 depletion, indicating that these components are particularly sensitive to the loss of TADA1-mediated protein-protein interactions, and that acute degradation of TADA1 might lead to collateral degradation of TAF12 due to the loss of its normal cotranslational assembly pathway. Although KAT2A is not known to directly interact with TADA1, its abundancy and engagement with SAGA depends on an intact structural CORE **(Fig. 1-3),** which becomes increasingly destabilised over time due to loss not only of TADA1, but the additional CORE subunits TAF12, TAF5L, and TAF6L (**Fig. 4E).** These results are therefore consistent with a cumulative disruption of the CORE-HAT interface following TADA1 depletion, which leads to HAT module disengagement and an increase in non-complexed KAT2A that becomes progressively destabilised in the absence of its normal binding partners. Altogether, these results indicate that acute perturbation of the SAGA CORE via depletion of TADA1 results in collateral degradation of selected SAGA components, particularly for proteins in the CORE and HAT modules that are structurally proximal to or direct interactors of TADA1.

### Non-complexed KAT2A is degraded in a process mediated by UBR5 and OTUD5

Perturbation of the SAGA CORE reduces KAT2A protein abundance while also resulting in disengagement of the HAT from the complex and the accumulation of low molecular weight KAT2A species. We therefore hypothesised that in CORE KO cells where complex integrity is perturbed, regions of KAT2A that in the intact SAGA complex are buried or bound by interaction partners might become exposed, resulting in the formation of degrons that target KAT2A to the proteasome and lead to its degradation. To assess if reduced KAT2A protein abundance in CORE KO mutants is indeed proteasome-sensitive, we therefore treated TAF5L KO HAP1 cells with inhibitors of key steps of the proteasome-mediated protein degradation machinery, including neddylation (MLN-4924), E1-enzyme activation (TAK-243), and the proteasome itself (MG-132), and measured endogenous KAT2A levels by immunofluorescence. Treatment with all three inhibitors restored KAT2A protein to wild type levels (**Fig. 5A, B**), with similar results in TAF6L and TADA1 KO cells **(Fig. S4A),** indicating a general proteasome-sensitivity of KAT2A when the SAGA CORE is perturbed. Crucially, in the absence of inhibitors, KAT2A mRNA levels remained unchanged or even increased in CORE KO cells compared to wild type (**Fig. S4B, C)**, demonstrating that loss of KAT2A protein is not due to reduced transcription. Thus, loss of KAT2A protein in the absence of *TAF5L, TAF6L,* or *TADA1* is the result of proteasome-mediated degradation.

**Figure 5.**
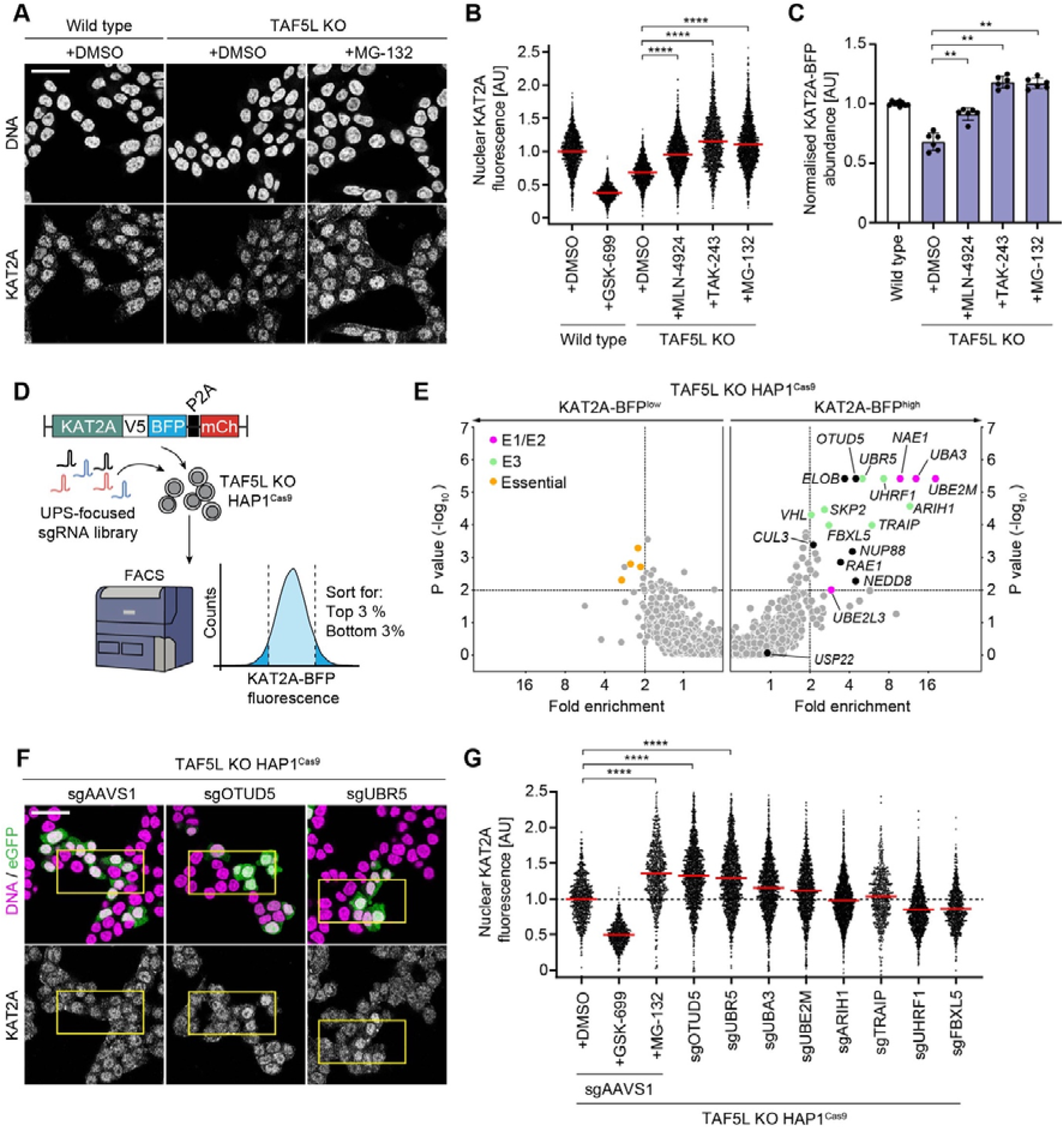
Non-complexed KAT2A is degraded in a process mediated by UBR5 and OTUD5. **A-C)** Inhibition of the proteasome, neddylation, or E1-activating enzymes rescues KAT2A protein levels in TAF5L KO cells. **A-B)** Wild type and TAF5L KO HAP1 cells were treated with DMSO, GSK-699 (100 nM), MLN-4924 (1 µM), TAK-243 (500 nM), or MG-132 (1 µM) for 8h as indicated, before fixation for immunofluorescence. **A)** Representative immunofluorescence images of wild type and TAF5L KO cells treated with compounds as indicated. Following compound treatment, cells were fixed and KAT2A was stained with an anti-KAT2A antibody. DNA was stained with Hoechst 33342. **(B)** Quantification of nuclear KAT2A fluorescence by immunofluorescence for wild type and TAF5L KO cells following compound treatment. Dots represent the mean nuclear KAT2A fluorescence of individual cells; red bars indicate the mean. Significance was tested using a two-tailed Mann–Whitney U-test. **C)** Quantification of normalised KAT2A-BFP fluorescence using the KAT2A stability reporter. Wild type and TAF5L KO cells expressing the KAT2A stability reporter were treated with inhibitors as in **A**, before analysis by flow cytometry. Dots represent the mean normalised KAT2A-BFP fluorescence of technical replicates, error bars represent the standard deviation, bars indicate the mean for each condition. Significance was tested using a two-tailed Mann–Whitney U-test. **D)** Schematic of FACS-based CRISPR–Cas9 screen to find regulators of KAT2A stability in TAF5L KO cells. TAF5L KO HAP1^Cas9^ cells expressing the KAT2A stability reporter were transduced with a UPS-focused gRNA library and sorted based on their KAT2A-BFP levels. **E)** Volcano plots of the FACS-based KAT2A stability reporter screen in TAF5L KO cells. Genes whose knockout reduces (left plot) or increases (right plot) KAT2A-BFP abundance are plotted. Fold changes and p-values of the KAT2A^high^ and KAT2A^low^ populations were calculated by comparison with the KAT2A^mid^ population using a two-sided negative binomial test (MAGeCK). Significant hits: Fold-enrichment ≥ 2 and -log_10_ p-values ≥ 2. Magenta dots represent significantly enriched E1/E2 enzymes. Green dots represent significantly enriched E3 substrate receptors. Orange dots represent essential genes. **F, G)** Validation of UPS effectors whose knockout increases KAT2A abundance by immunofluorescence. Top hits that increased KAT2A abundance in the pooled screen were validated by transduction of TAF5L KO^Cas9^ HAP1 cells with GFP expressing gRNAs, with the two top-scoring gRNAs transduced independently for each UPS gene. 5 days post-transduction, cells were fixed and KAT2A was stained with an anti-KAT2A antibody. DNA was stained with Hoechst 33342. Where indicated, cells were treated with GSK-699 (100 nM) or MG-132 (1 µM) for 8 h before fixation. All other samples were treated with DMSO. **F)** Representative immunofluorescence images of TAF5L KO HAP1^Cas9^ cells transduced with gRNAs against *AAVS1*, *OTUD5*, or *UBR5*, as indicated. **G)** Quantification of nuclear KAT2A fluorescence of TAF5L KO HAP1^Cas9^ cells transduced with the indicated gRNAs. GFP positive cells were analysed. The data for two independent gRNAs is merged for each UPS gene. Dots represent the mean nuclear KAT2A fluorescence of individual cells; red bars indicate the mean. Significance was tested using a two-tailed Mann–Whitney U-test. Data information: **B, G** (****) P < 0.0001; two-tailed Mann–Whitney U-test. **C** (**) P < 0.01; two-tailed Mann–Whitney U-test. Biologically independent replicates: **(A-C, F, G** (n = 2), **E** (n = 3). In **B** the data was normalised relative to the mean nuclear KAT2A fluorescence of wild type HAP1 cells. In **G** only GFP positive cells were analysed, and data was normalised relative to the mean nuclear KAT2A fluorescence of AAVS1 transduced TAF5L KO HAP1^Cas9^ cells treated with GSK-699. All images show single Z-slices. Yellow boxes indicate inset regions showing non-transduced (GFP negative) and transduced (GFP positive) cells. Scale bars: 20 µm. The number of cells analysed by microscopy in **B, G** is listed in **Table S3.**

Having determined that the loss of KAT2A protein is proteasome-sensitive, we wished to understand the molecular mechanism of KAT2A degradation in CORE KO mutants and identify which of the effectors of the Ubiquitin-proteasome system (UPS), including the more than 600 E3 ligases in the human genome, targets KAT2A for degradation. To do so, we again made of use of the KAT2A stability reporter **(Fig. 1).** Treatment of TAF5L KO or TAF6L KO cells expressing the stability reporter with inhibitors of the proteasome machinery also rescued KAT2A-BFP protein levels (**Fig. 5C, S4D)**, thus allowing screening for effectors of KAT2A degradation. To identify specific effectors regulating KAT2A protein stability, we therefore performed a FACS-based pooled CRISPR screen in TAF5L KO HAP1^Cas9^ cells, with a library targeting 1301 Ubiquitin-associated human genes, with 6 gRNAs per gene (Kagiou *et al*, 2024) (**Fig. 5D**). By sorting cells expressing high levels of KAT2A BFP (KAT2A-BFP^high^) after library transduction (**Fig. S4E)** and performing next-generation sequencing (NGS), we could identify genes whose knockout promotes KAT2A stabilisation to nominate candidate regulators of KAT2A degradation when the SAGA CORE is perturbed.

Interestingly, knockout of USP22, which confers the enzymatic activity of the SAGA DUB module and could therefore in principle auto-deubiquitinate KAT2A, did not alter KAT2A abundance (**Fig. 5E)**, ruling out the possibility of interactions between the HAT and DUB modules regulating KAT2A stability. Consistent with experiments with inhibitors, loss of components of the neddylation machinery, which are required for activation of Cullin Ring E3 Ligases (CRLs), scored as top hits in the screen, including the E1-enzymes *NAE1* and *UBA3*, which form a heterodimer, the E2-conjugating enzyme *UBE2M,* and *NEDD8*, which encodes the neddylation modification itself (**Fig. 5E**). However, despite this neddylation sensitivity, we found only weak enrichment of CRL components, such as the E2 ubiquitin-conjugating enzyme *UBE2L3,* the E3 ligase substrate receptor *VHL,* and the E3 scaffolding protein *CUL3,* suggesting non-specific or promiscuous degradation of KAT2A by CRLs that is abrogated only upon inhibition of all of them, which would occur upon loss of neddylation pathway components (**Fig 5E)**. We however uncovered numerous candidate non-CRL E3 ligases that were significantly enriched in KAT2A-BFP^high^ cells, such as the DNA-repair associated *TRAIP* (Harley *et al*, 2016), the RBR (RING-between-RING) E3 ligase *ARIH1,* and *UHRF1* (Vaughan *et al*, 2018), a known chromatin-associated factor. We also identified the HECT-type E3 ligase *UBR5* (Mansfield *et al*, 1994; Hodáková *et al*, 2023; Wang *et al*, 2023), in addition to its interactor, the deubiquitinase *OTUD5* (de Vivo *et al*, 2019), as significant hits whose loss stabilised KAT2A in the absence of TAF5L (**Fig. 5E**).

To validate which of the candidate UPS effectors promote KAT2A degradation upon SAGA CORE perturbation, we knocked out the top hits from the pooled screen with the two top-scoring guide RNAs for each gene in TAF5L KO HAP1^Cas9^ cells, and used confocal microscopy to measure endogenous KAT2A protein levels, using GFP as a marker to identify transduced cells. To better assess the relative rescue of KAT2A protein levels in CORE KO cells, wild type HAP1^Cas9^ cells were transduced with the same guides, with cells treated with GSK-699 or MG-132 serving as normalisation controls to determine the dynamic range of the assay. In agreement with findings from the pooled CRISPR screen, knockout of the neddylation pathway components *UBE2M* and *UBA3* rescued KAT2A protein levels in TAF5L KO cells **(Fig. 5F, G),** although only partially, reaching approximately 50 % of the levels observed upon proteasome inhibition. Loss of neddylation components was however as effective as proteasome inhibition in wild type cells, with a mild but significant increase in nuclear KAT2A signal **(Fig. S4F, G**), indicative of a reduced contribution of neddylation-dependent E3 ligases to KAT2A protein degradation when the SAGA CORE is perturbed. Interestingly, the magnitude of rescue of KAT2A protein levels upon proteasome inhibition was substantially higher in TAF5L KO cells compared to wild type (**Fig. 5F, G, S4F, G),** with a greater than two-fold relative increase in mean KAT2A fluorescence (normalised mean KAT2A fluorescence: TAF5L KO sgAAVS1 + MG-132, 1.35 ± 0.42; Wild type sgAAVS1 + MG-132, 1.16 ± 0.33, mean + S.D.). Similar results were also observed in TADA1 KO cells **(Fig. S4H-I),** indicating a general increase in the sensitivity of KAT2A to proteasome-mediated degradation when the CORE is perturbed. Such findings are consistent with an increased abundance of low molecular weight KAT2A species in CORE KO mutants (**Fig. 2D, E),** where a greater proportion of KAT2A likely exists outside of the intact SAGA complex. Loss of binding interactions with other SAGA components upon SAGA CORE disruption might therefore render KAT2A more susceptible to collateral protein degradation when complex integrity is disrupted, and thus more responsive to stabilisation upon proteasome inhibition compared to wild type cells, where the HAT module remains engaged with SAGA.

Most strikingly, knockout of both *UBR5* and *OTUD5* rescued nuclear KAT2A protein levels to the same extent as proteasome inhibition in both TAF5L KO and TADA1 KO cells. In contrast, the magnitude of rescue was more than two-fold lower in wild type cells (**Fig. 5F, G, S4F-I**), implicating UBR5 and OTUD5 as specific regulators of KAT2A degradation when the SAGA CORE is perturbed. Consistent with *UBR5*-mediated KAT2A degradation, OTUD5 is known to stabilise *UBR5* via lysine deubiquitination, with loss of *OTUD5* thus also reducing UBR5 protein levels (de Vivo *et al*, 2019). Intriguingly, *UBR5* was recently identified as a key regulator of homeostatic protein turnover, functioning as a E3 ligase dedicated to the degradation of solitary or orphan proteins, including transcription factors such as *MYC,* and nuclear hormone receptors (Schukur *et al*, 2020; Tsai *et al*, 2023; Mark *et al*, 2023), indicating that UBR5 might specifically recognise non-complexed KAT2A as an orphan substrate when the SAGA CORE is disrupted, leading to its collateral degradation.

Taken together, we therefore propose a working model in which the structural integrity of the SAGA complex is dependent on its non-enzymatic CORE subunits, TAF5L, TAF6L, and TADA1, with complex integrity essential for engagement of the HAT module with SAGA, and consequently for the stability and function of its histone acetyltransferase, KAT2A (**Fig. 6A**). Disruption of the SAGA CORE leads to disengagement of the HAT from SAGA, potentially via disruption of co-translational or early post-translational assembly, and leads to the accumulation of solitary, unassembled KAT2A protein that is not recruited to chromatin and is instead targeted for proteasomal degradation. This degradation is mediated, at least in part, by orphan quality control factors such as the E3 ligase UBR5, in concert with the deubiquitinase OTUD5, which is known to regulate UBR5 stability **(Fig. 6A**). Thus, the stability and activity of KAT2A is tightly coupled to the structural organisation of the SAGA complex, and perturbing its assembly triggers a quality control response that selectively eliminates unassembled KAT2A, with functional consequences for histone acetylation and gene regulation. More broadly, our study highlights a general vulnerability in multi-subunit co-activator complexes that could be exploited therapeutically to target chromatin-modifying enzymes in cancer. By targeting structural complex components that themselves lack enzymatic activity, it is possible to disrupt complex function through two distinct yet complementary mechanisms as: 1) loss of complex integrity impairs recruitment of enzymatic subunits to chromatin, leading to reduced catalytic output due to loss of interactions between the fully assembled complex and its substrate and 2) leads to increased incidence of orphan complex subunits that can become collaterally degraded in the absence of the intact complex, further reducing complex functionality.

**Figure 6.**
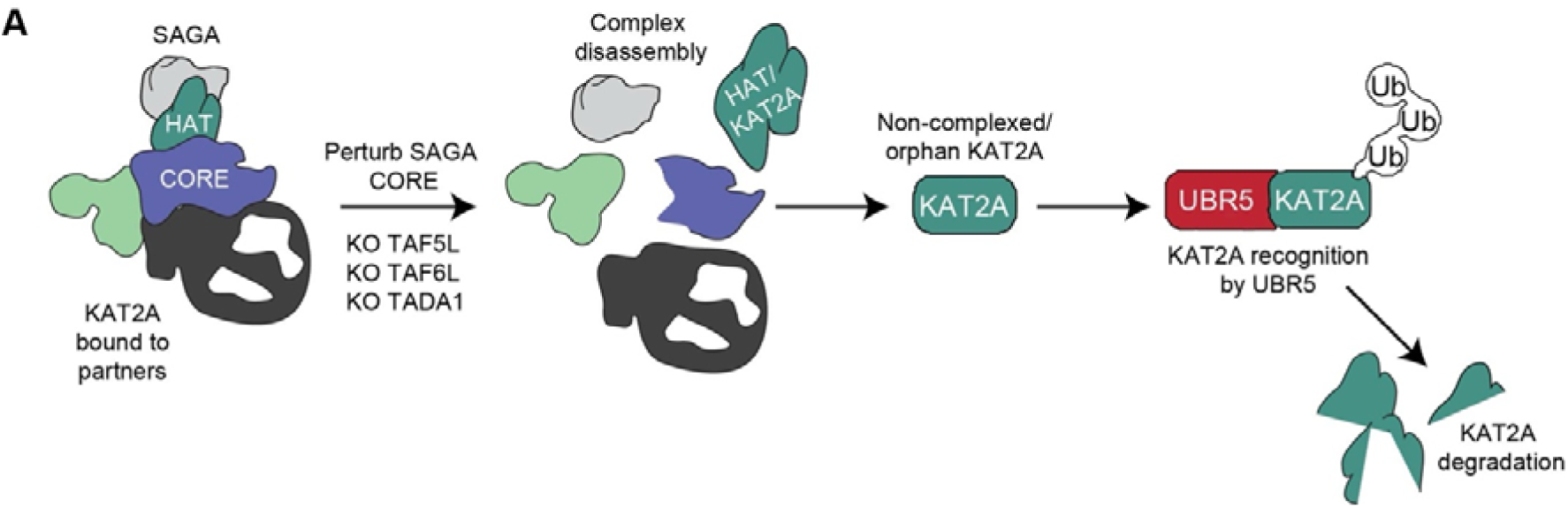
Model of KAT2A degradation following disruption of the SAGA CORE. **A)** Loss of the SAGA CORE components TAF5L, TAF6L, or TADA1, results in disassembly of the SAGA complex. Complex disassembly leads to increased abundance of free floating KAT2A protein, converting it into a substrate that is recognised by the E3 ligase UBR5, thereby funnelling KAT2A into the orphan protein degradation pathway.

## DISCUSSION

Our study identifies a mechanism by which the structural integrity of the SAGA co-activator complex regulates the stability and function of one of its catalytic subunits, KAT2A. We demonstrate that a subset of non-enzymatic components of the SAGA CORE module, namely TADA1, TAF5L, and TAF6L, are required to maintain KAT2A protein abundance and activity (**Fig. 1**). Loss of these structural subunits leads to disengagement of the HAT module, mislocalisation of KAT2A from chromatin (**Fig. 2, 3**), and its subsequent degradation via the ubiquitin-proteasome system (**Fig. 5**) with concurrent, collateral degradation of other elements of the CORE and HAT modules (**Fig. 4**). These findings reveal that SAGA-dependent gene regulation relies not only on the enzymatic activities of its HAT and DUB modules but also on a finely tuned assembly architecture that protects vulnerable catalytic components from orphan degradation.

Our work builds on the emerging paradigm that many multi-subunit protein complexes are assembled co-translationally and are dependent on precise stoichiometric balances to prevent the accumulation of orphan proteins (Natan *et al*, 2017; Bernardini & Tora, 2024). By perturbing specific SAGA CORE components, we effectively uncouple KAT2A from its native assembly pathway, rendering it susceptible to proteasomal clearance. The gradual depletion of KAT2A following acute degradation of TADA1 suggests that CORE components may serve a chaperone-like function during or shortly after KAT2A translation, ensuring its successful incorporation into the SAGA complex. This process extends to other subunits that are directly interacting with TADA1, such as its co-translational assembly partner TAF12 (Xu *et al*, 2018; Yayli *et al*, 2023), or other subunits in close proximity to TADA1 in the CORE and HAT modules, including TAF5L, TAF6L, and TADA3 (Herbst *et al*, 2021). Altogether these results show progressive disruption of SAGA complex integrity and HAT module engagement upon CORE perturbation, ultimately leading to an increased abundance of non-complexed KAT2A that becomes a substrate for collateral protein degradation.

Importantly, we identify UBR5 and OTUD5, previously linked to orphan protein quality control and homeostatic turnover of key transcription factors such as MYC (Schukur *et al*, 2020; Mark *et al*, 2023), as critical effectors of collateral KAT2A degradation. These findings suggest that once SAGA disassembles, exposed degrons or misfolded regions within KAT2A are recognised by the orphan surveillance machinery, tagging the protein for proteasomal degradation. Notably, inhibition of key proteasomal regulators or genetic disruption of UBR5 and OTUD5 restores KAT2A protein levels, underscoring the specificity of this degradation pathway.

This work has broader implications for therapeutic strategies targeting multi-protein complexes in cancer. While much attention has focused on inhibiting enzymatic activities within multisubunit protein complexes such as SAGA, our data provides further evidence that destabilising critical structural subunits may offer an alternative approach, as targeting non-enzymatic scaffolding proteins could lead to secondary degradation of catalytic components, amplifying the therapeutic impact. Such a strategy could be particularly attractive in cancers dependent on SAGA-mediated acetylation programs, as KAT2A has been identified as a cancer vulnerability in multiple contexts (Tzelepis *et al*, 2016; Malone *et al*, 2024). Indeed, collateral degradation of SAGA components could contribute to the dependencies observed in neuroblastoma and leukemia (Barbosa *et al*, 2024; Malone *et al*, 2024).

In this context, small molecules targeting TAF5L, TAF6L or TADA1 would abrogate KAT2A function on two fronts: first, via proteasome-mediated collateral degradation, reducing KAT2A protein abundance, and secondly, by disrupting complex integrity and chromatin binding. The latter is likely to affect not only KAT2A, but also its paralogue KAT2B, which although not the predominant acetyltransferase found within SAGA, can be incorporated into SAGA instead of KAT2A in a mutually exclusive fashion (Nagy & Tora, 2007). Given the functional redundancy between KAT2A and KAT2B (Malone *et al*, 2024), targeting shared structural scaffold subunits therefore has the potential to simultaneously target both paralogues at the same time, an approach that could be exploited therapeutically in KAT2A-dependent cancers (Tzelepis *et al*, 2016; Malone *et al*, 2024), in addition to or in combination with KAT2A/B degraders that have been developed (Bassi *et al*, 2018; Malone *et al*, 2024; Liu *et al*, 2025), but not yet tested in patients.

Future work will be necessary to dissect how broadly this mechanism applies to other multi-subunit transcriptional regulators and to fully define the degrons within KAT2A that trigger its orphan degradation. It will also be important to determine whether additional components of the orphan quality control machinery are selectively engaged depending on the nature of the disrupted complex, and whether those can be specifically hijacked using proximity-inducing pharmacology. Given the essential role of transcriptional co-activators like SAGA in cell fate regulation, these insights could pave the way for novel approaches to target transcriptional addiction in cancer and other diseases.

## METHODS

### Cell lines

All HAP1 cell lines used in this study are derived from a parental HAP1 cell line (Horizon Discovery C631). HAP1 cells were cultured in IMDM (Gibco) supplemented with 10 % (v/v) fetal bovine serum (Gibco, 10270-106), 1 % (v/v) Penicillin–Streptomycin (Fisher Scientific, 11548876), L-Glutamine (Thermo Fisher, 25030024). HEK293T cells were cultured in DMEM supplemented as for IMDM. Cells expressing Blasticidin or Puromycin resistance cassettes were cultured in medium supplemented with either 10 µg ml^-1^ Blasticidin (Gibco, A1113903) or 1 µg ml^-1^ Puromycin (Thermo Fisher Scientific, A1113803). All cell lines were cultured in a humidified incubator at 37°C with 5 % CO_2_, and regularly tested negative for mycoplasma contamination.

### Lentivirus production

HEK293T cells were seeded into 15 cm dishes ∼ 24 hours prior to transfection. Cells were transfected at 80 % confluence in 16 ml of media with 8.75 μg of VSV-G, 16.25 μg of psPAX2, and 25 μg of the CRISPR-Cas9 pooled lentiviral plasmids, using 150 μg of linear polyethylenimine (PEI) (Sigma-Aldrich). Media was changed with fresh media 16–24 hours post-transfection. Lentiviral supernatant was collected 48 and 72 hours post-transfection and subsequently concentrated by ultracentrifugation (24000 rpm, 4°C, 2 hours). For crude lentiviral supernatants, transfections were performed in a 6-well format. Briefly, cells were transfected at 80 % confluence in 2 ml of media with 350 ng of VSV-G, 650 ng of psPAX2, and 1 μg of the lentiviral construct using 6 μg of PEI. Media was exchanged 16–24 hours post-transfection. The supernatant was collected 48 and 72 hours post-transfection as separate harvests and centrifuged to remove cellular debris (4700 rpm, 4°C, 10 min). Crude lentiviral supernatant was diluted with complete medium before transducing cells, as specified below.

### Generation of HAP1^Cas9^ cells

To generate HAP1 cells stably expressing Cas9 in both wild type and TAF5L KO backgrounds, cells were transduced with Cas9-Blast lentivirus (generated from pLentiCas9-Blast, Addgene, 52962) and selected with 10 µg ml^-1^ Blasticidin (Gibco, A1113903) for 7 days. Clones were obtained by limited dilution into 96-well plates. Cas9 activity was determined with a competition assay using gRNA targeting either the essential gene *RAD21* or the safe harbour locus *AAVS1,* cloned into a GFP-expressing vector (pLKO5.gRNA.EFS.GFP (Addgene, 57822)). GFP fluorescence was measured 3-and 7-days post-transduction by flow cytometry using an LSR Fortessa instrument (BD Biosciences) with an HTS Loader. Clones showing dropout of GFP fluorescence upon targeting *RAD21* but not *AAVS1* were selected.

### Generation of HAP1 knockout clones

gRNAs were designed to target critical exons of the selected candidate genes and cloned into pLKO5.gRNA.EFS.GFP. One day before transfection 400,000 HAP1^Cas9^ cells were seeded into 6-well plates. The following day cells were transfected with 2 µg gRNA plasmid in total (1 µg per gRNA plasmid if using multiple plasmids) using Turbofect Transfection Reagent (Thermo Fisher Scientific, R0531). 2 – 3 days post transfection GFP positive cells were sorted using a FACS Aria Fusion instrument (BD Biosciences) and replated into 6-well plates. Cells were allowed to recover in fresh complete medium for one day. The following day cells were detached from the culture plate and seeded at clonal density into 96-well plates. Following clonal expansion, individual clones were genotyped by PCR and Sanger sequencing to detect the homozygous/biallelic deletion or *knockout* (KO) clones. Primers used for genotyping are listed in **Table S1**.

### Generation of KAT2A stability reporter expressing cells

The KAT2A stability reporter was generated by modifying a previously published BRD4 reporter plasmid (Hsia *et al*, 2024). The BRD4 cDNA sequence was excised and replaced with that of KAT2A by SalI/BamHI digestion and Gibson Assembly. The KAT2A ORF was PCR-amplified from TFORF2959 (Addgene, 144435). The resultant pRRL-SFFV-KAT2A-GGGG-3xV5-TagBFP-P2A-mCherry plasmid was produced as crude lentivirus as described above and used to transduce HAP1 cells. 7 - 10 days post transduction, cells were sorted for mCherry positive cells using a FACS Aria Fusion instrument (BD Biosciences). Cells expressing low, medium, and high levels of mCherry were sorted. For the arrayed loss of function CRISPR screen performed in wild type HAP1^Cas9^ cells, cells expressing high levels of mCherry were used. For the pooled gain of function CRISPR screen performed in TAF5L KO^Cas9^ cells and validation experiments using inhibitors, cells expressing low to medium levels of mCherry were used.

### Arrayed CRISPR screening using the KAT2A stability reporter

Crude lentiviral supernatants for gRNAs were generated as described above and distributed in an arrayed format in 96-well plates. One day prior to transduction, wild type HAP1^Cas9^ cells expressing the KAT2A stability reporter, in addition to the parental cell line without the reporter, were seeded into F-bottomed 96-well plates (VWR, 734-2327) at 7,000 cells per well in complete IMDM. For seeding, cells were washed once with PBS and incubated with Accutase® Cell Detachment Solution (Thermo Fisher, 423201) for 3-5 min at room temperature, quenched with complete medium, and filtered once through a 35 µm nylon mesh before counting. On the day of transduction, cells were transduced with 100 µl of crude viral supernatant (50 µl of crude supernatant diluted 1:1 with complete medium), with one guide RNA transduced per well (in triplicate). Both the parental cell line and reporter expressing cells were transduced with each guide RNA in the arrayed collection. 3, 5, and 7 days after transduction, half of the cells in each well were analysed by flow cytometry. The remaining half were passaged, with fresh complete medium added to a final volume of 200 µl per well. For flow cytometry, the medium was removed from each well, cells washed twice with 100 µl PBS, and incubated with 100 µl Accutase for 5 – 10 min at room temperature. Cells were resuspended in the plate and 50 µl of the cell suspension transferred to a V-bottomed 96-well plate (Thermo Fisher, 277143) containing 50 µl of FACS buffer (PBS containing 2 % FCS, 2 mM EDTA pH 8.0 (Thermo Fisher, AM9260G)). BFP, GFP and mCherry fluorescence was measured for each well using an LSR Fortessa instrument (BD Biosciences) with attached HTS Loader. Flow cytometry data was analysed using FlowJo V10.10.0. To calculate the gRNA dropout for each gene, the fraction of GFP positive cells was determined for each guide at each time point. The data was normalised independently for each guide relative to the fraction of GFP positive cells 3 days after transduction. To calculate KAT2A-BFP protein abundance 5 days after lentiviral transduction, cells were first gated for GFP positive cells (to select only transduced cells) and then mCherry positive cells (to select only cells expressing the reporter). To account for different transduction efficiencies between different GFP expressing gRNAs, BFP and mCherry mean intensity values were normalised by background subtraction of the respective values for reporter negative cells transduced with the same gRNA. Relative KAT2A abundance was calculated as the ratio of background subtracted BFP to mCherry mean fluorescence intensity for each gRNA. Wild type HAP1^Cas9^ cells expressing the KAT2A stability reporter were gated in the following way: 1) FSC-A vs SSC-A to gate cells, 2) FSC-A vs FSC-H and SSC-H vs SSC-W to gate single cells, FSC-A vs GFP-A to gate GFP positive cells, and then KAT2A-BFP-A vs mCherry-A to measure the KAT2A-BFP/mCherry ratio. For stability reporter experiments in **Fig. 5C**, **S4D**, the same gating strategy was used, with the exception of gating for GFP positive cells as the experiment was performed on GFP negative cells. The data in **Fig. 1C** is plotted relative to the mean KAT2A abundance of sgAAVS1 transduced reporter cells treated with the KAT2A/B PROTAC GSK-699 (Malone et al, 2024) (0 value) and the mean KAT2A abundance of the three negative control guide RNAs from the arrayed collection (1 value). GSK-699 was used at a concentration of 100 nM for 6 h. A list of the plasmids used and the sequences of the gRNAs can be found in **Table S1.**

### Pooled CRISPR screen using the KAT2A stability reporter

A FACS-based CRISPR/Cas9 screen using KAT2A stability reporter cells was performed, followed by library preparation, NGS, and data analysis, as previously described with minor modifications (de Almeida *et al*, 2021; Hsia *et al*, 2024; Kagiou *et al*, 2024). Briefly, a Ubiquitin–proteasome system (UPS)-focused sgRNA library targeting 1301 human ubiquitin-related genes with 6 sgRNAs per gene (Kagiou *et al*, 2024) was used to transduce TAF5L KO HAP1^Cas9^ KAT2A-BFP-P2A-mCherry cells. Transductions were performed at a multiplicity of infection (MOI) of 0.1, ensuring a 1000-fold library coverage. Ten days after G418 selection (1CmgCml⁻¹; Sigma-Aldrich, A1720), 50 million cells per replicate were harvested (three replicates harvested). Following selection, cells were incubated for 10 min at 4°C with an anti-Thy1.1–APC antibody (BioLegend, 202526; 1:400) and Human TruStain FcX™ Fc Receptor Blocking Solution (BioLegend, 422301; 1:400). Cells were fixed with 4% BD Fixation Buffer (Thermo Scientific™ Pierce™, BD 554655) for 45 min at 4 °C, protected from light, and stored overnight at 4°C in PBS containing 5% FBS and 1 mM EDTA. The next day, cells were sorted on a BD FACSAria™ Fusion (BD Biosciences) using a 70 μm nozzle. Aggregates, dead cells, and sgRNA library–negative (Thy1.1–APC–negative) cells were excluded. The remaining population was sorted into KAT2A^high^ (∼3% of cells), KAT2A^mid^ (∼30%) and KAT2A^low^ (∼3%) fractions based on KAT2A-BFP and mCherry expression, ensuring a minimum of 1500-fold library coverage per replicate. Genomic DNA from sorted cell populations was extracted by cell lysis (10 mM Tris-HCl, 150 mM NaCl, 10 mM EDTA, 0.1% SDS), followed by proteinase K digestion (New England Biolabs, P8107) and RNA removal using DNase-free RNase A (Thermo Fisher Scientific, EN0531). DNA was purified by two rounds of phenol extraction (Sigma-Aldrich, P4557) and isopropanol precipitation (Sigma-Aldrich, I9516). The gRNA cassettes were then amplified in two PCR steps using AmpliTaq Gold polymerase (Thermo Fisher Scientific, 4311818), with sample-specific barcodes incorporated in the first PCR and Illumina adapters added in the second. PCR products were cleaned using Mag-Bind TotalPure NGS beads (Omega Bio-tek, M1378-00), pooled, and sequenced on an Illumina NovaSeq 6000 platform. The screen data was processed using the crispr-process-nf Nextflow pipeline (https://github.com/ZuberLab/crispr-process-nf/). In brief, raw FASTQ files were trimmed with *cutadapt* (v4.4) to remove random barcodes and spacer sequences, followed by demultiplexing based on sample barcodes. Reads were aligned to the custom UPS sgRNA reference library using *Bowtie2* (v2.4.5), and sgRNA abundance was quantified with *featureCounts* (v2.0.1). The resultant count tables (**Table S2**) were analysed using the crispr-mageck-nf workflow (https://github.com/ZuberLab/crispr-mageck-nf/) for downstream statistical evaluation. Gene-level enrichment was assessed using *MAGeCK* (v0.5.9) (Li et al., 2014), comparing KAT2A^high^ or KAT2A^low^ populations to the KAT2A^mid^ reference group, based on median-normalised read counts and replicate-level variance estimation.

### Immunofluorescence

For cell seeding, cells were washed once with PBS and incubated with Accutase for 3-5 min at room temperature, quenched with complete medium and filtered once through a 35 µm nylon mesh, before cell counting. For confocal microscopy experiments, cells were seeded at 50,0000 cells per well one day prior to fixation in 8-well Ibidi chambers (Ibidi, 80827). For arrayed CRISPR screening experiments in 96-well plates, 7000 - 10000 cells were seeded per well one day prior to transduction in F-bottomed 96-well plates (VWR, 734-2327). The following day, cells were transduced with 100 – 150 µl of crude viral supernatant (crude viral supernatant diluted 1:1 with complete medium). 3 days after transduction lentiviral medium was removed, the cells detached from the plate using Accutase and then transferred to imaging plates (PhenoPlate 96-well plates (black, optically clear flat-bottom, tissue-culture treated (Revvity, 6055302)) or 8-well Ibidi chambers. Cells were fixed for immunofluorescence 5 days (KAT2A staining) or 6 days (H3K9ac staining) post-lentiviral transduction. At the time of fixation, cells were washed twice with PBS before fixation with PBS containing 4 % methanol-free formaldehyde (Thermo Fisher Scientific, 28906) for 5 min (Ibidi chambers) or 15 min (96-well plates). The fixative was removed and cells quenched using 10 mM Tris–HCl (Invitrogen, 15567027) pH 7.5 in PBS for 3 min. For staining of nuclear antigens, cells were permeabilised with PBS containing 0.2 % Triton-X-100 (Sigma Aldrich, X100-100 ml) for 5 min or 0.5 % Triton X-100 for 15 min (H3K9ac staining experiments only), washed once with PBS to remove residual detergent, and blocked for 30 min with 0.45 µm filtered 2% BSA (Sigma-Aldrich, A9418-50G) in PBS (blocking buffer). Primary and secondary antibodies were diluted with blocking buffer. Primary antibody incubations were performed for > 16 h at 4°C with gentle shaking, followed by 3 × 10 min washes with PBS, and incubation with secondary antibodies for 1 h at room temperature with gentle shaking. Samples were then washed 3 × 10 min with PBS containing 1.62 µM Hoechst 33342 (Thermo Fisher Scientific, H3570). Antibodies used for immunofluorescence and their working dilutions can be found in the **Key Resources Table**.

### Microscopy and image processing

All confocal microscopy experiments, with the exception of **Fig 1.D, E,** were imaged on a custom Zeiss LSM 980 microscope fitted with an additional Airyscan2 detector, using a 63× NA 1.4 oil DIC Plan-Aphrochromat (Zeiss) objective and ZEN 3.3 Blue 2020 software. The data in experiment **1D, E**, was acquired on an Opera Phenix high-content confocal microscope (PerkinElmer) using a 20x air objective and imaging 18 fields of view per well. For images acquired using the Opera Phenix system, individual channels were flat-field corrected using Python following acquisition. The flat-field corrected images were then merged into multichannel composite images, which were used as the input for the analysis pipeline.

### Image analysis – measurement of mean pixel intensities in fields of cells

Fields of cells were analysed using a custom automated analysis pipeline written in Python. Before measuring mean nuclear fluorescence of markers of interest, the following criterion was pre-established: only interphase cells were considered for the analysis. For most immunofluorescence experiments, H3S10P was also stained as a mitotic marker, with H3S10P positive cells excluded from the analysis. Size and maximum mean DNA fluorescence filters were applied to the data to also help exclude mitotic cells. A Gaussian blur was applied to the Hoechst channel using the skimage ‘filters’ module and the Hoechst channel was then segmented using the ‘nuclei’ module of cellpose (Stringer et al, 2021). Cells which touched the border of the image were excluded using the ‘clear_border’ functionality from the skimage ‘segmentation’ module. Individual masks were labelled and applied to each cell in the field. The area of the nuclear mask and mean fluorescence within the nuclear mask was then calculated for each segmented cell, and the data output as a csv file for each input condition. Biological replicates were normalised independently, relative to their own internal positive and negative controls and the normalised data then merged for the final figure. To account for differences in cell ploidy and the local compaction state of the chromatin, in **Fig. 2D** the H3K9ac signal level for each condition was determined by dividing the mean nuclear H3K9ac fluorescence of each segmented nucleus by the mean fluorescence of the DNA (stained with Hoechst 33342), to calculate a H3K9ac/Hoechst ratio.

### RNA isolation and RT-qPCR

RNA was isolated from cells using a RNeasy Mini Kit (Qiagen, 74104) according to the manufacturer’s instructions. 500 ng RNA was used for cDNA preparation with an iScript cDNA Synthesis Kit (Bio-Rad, 170889). RT-qPCR was performed using an iQ SYBR Green Supermix (Bio-Rad, 1708880) on a Bio-Rad iCycler RT-PCR detection system. Primers used for RT-qPCR are listed in **Table S1.**

### CUT&RUN sample preparation

For each sample, nuclei from 250,000 cells were isolated in 100 µl nuclear extraction (NE) buffer (20 mM HEPES (Thermo Fisher, 15630-056), 10 mM KCl (Thermo Scientific, AM960-G), 0.1 % Triton X-100 (Sigma, 93443-100ML), 20 % Glycerol (Fisher Scientific, 10692372), 0.5 mM Spermidine (Sigma, S2626-1G) supplemented with Roche cOmplete EDTA-free Protease Inhibitors (Sigma, 11873580001) for 10 min on ice, centrifuged for 3 min at 600 g at room temperature, resuspended in 100 µl NE buffer and bound to 10 µl of activated ConA beads for 10 min at room temperature. Next, beads were collected with a magnetic rack and resuspended in 50 µl Antibody Buffer (20 mM HEPES, 150 mM NaCl (Fisher Scientific, BP358-1), 0.5 mM Spermidine, 0.02 % Digitonin (Millipore, D141-100MG), 2 mM EDTA supplemented with Roche Complete EDTA-free Protease Inhibitors) supplemented with 1 µl of antibody. Nuclei were incubated overnight on a nutator at 4C. After two washes with 250 µl Digitonin buffer (20 mM HEPES, 150 mM NaCl, 0.5 mM Spermidine, 0.02 % Digitonin supplemented with Roche Complete EDTA-free Proteasome Inhibitor), nuclei were resuspended in 50 µl Digitonin buffer supplemented with 700 ng/µl pAG-Mnase (gift of Joris Van der Veeken, IMP, Vienna), and incubated for 30 min at room temperature. Excess pAG-Mnase was removed with two washes with Digitonin buffer. Finally, nuclei were resuspended in 50 µl Digitonin buffer and pAG-Mnase was activated with 1 µl 100 mM CaCl_2_ for 1.5 h on ice. The reaction was quenched with 33 µl Stop Buffer (340 mM NaCl, 20 mM EDTA, 4 mM EGTA (VWR, J60767AD), 50 µg/mL RNAse A, 50 µg/mL Glycogen (Merck, 10901393001). Cleaved DNA was allowed to diffuse out by incubating the tubes at 37°C for 10 min. After separation using a magnetic rack, DNA was isolated using the Monarch DNA Cleanup Kit (NEB, T1030) and eluted in 15 µl Elution Buffer. Next-generation sequencing libraries were prepared from 2 ng DNA using the NEBaNext Ultra II DNA Library prep Kit (NEB, E7645L), pooled and sequenced (PE150) at Azenta, at a depth of 5-10M reads per library. Antibodies used for CUT&RUN and their working dilutions can be found in **Key Resources Table.**

### CUT&RUN data analysis

Sequencing reads (150bp paired-end, NovaSeq 6000) were trimmed using TrimGalore and aligned to the human genome, hg38, using Bowtie2 (Langmead & Salzberg, 2012) (--local, -- very-sensitive, --no-mixed, --no-discordant, --dovetail). SAMtools (Li *et al*, 2009) was used to fix mates of paired end reads, merge, sort and index bam files, while a CUT&RUN-specific blacklist (Nordin *et al*, 2023) was removed from the bam files using BEDtools (Quinlan & Hall, 2010). DeepTools (Ramírez *et al*, 2016) was then used to build the genome coverage, to compute matrices and plot the profiles of KAT2A aligned around transcriptional start sites.

### Sucrose gradient fractionation

Sucrose (Fisher Scientific, 15446759) stock solutions (10 % and 60 %, prepared in dH_2_O), were diluted 1:1 with 2X NP40-Lysis buffer (40 mM Tris HCl pH 7.5 (Invitrogen, 15567027), 200 mM NaCl (Fisher Scientific, BP358-1), 10 mM MgCl_2_ (Merck, MG1028-100ML), 0.4 % NP40 (Thermo Fisher, 85124) to generate 5 and 30 % sucrose solutions in 1X NP40-Lysis buffer. DTT (Sigma, 43816-10ML) was added to a concentration of 1 mM to diluted sucrose solutions immediately before gradient preparation. 5 – 30 % sucrose gradients were prepared in ultracentrifuge tubes (Open-Top Polyclear Centrifuge Tubes, 14 x 89 mm, Seton Scientific, 7030) using a BioComp gradient station (Programme: Long cap, Sucrose, 05-30 %, weight/volume), and subsequently equilibrated to 4°C for > 1 h before sample loading. The rotor was also equilibrated to 4°C before ultracentrifugation. Samples were harvested freshly before ultracentrifugation. Cells growing in 15 cm plates were harvested by washing once with ice-cold PBS, with banging against a solid support to remove dead cells. PBS was removed and cells were detached by incubation with Accutase for 3 - 5 min at room temperature. Cells were then quenched with complete medium, centrifuged (400 g, 5 min, 4°C), washed once with ice-cold PBS, centrifuged once more and transferred to 1.5 ml Eppendorf tubes in 1 ml ice-cold PBS. Cells were centrifuged (1100 g, 1 min, 4°C), PBS removed, and cell pellets lysed in 150 µl 1X NP40-lysis buffer (20 mM Tris HCl pH 7.5, 100 mM NaCl, 5 mM MgCl_2,_ 0.2 % NP40, 1 mM DTT, 2X Roche cOmplete EDTA-free protease inhibitors (Sigma, 11873580001)) for 30 min on ice, with vortexing every 10 min. Lysates were clarified by centrifugation (17000 g, 15 min, 4°C) and the concentration of the clarified supernatant was quantified using a BCA kit (Pierce, Thermo Scientific, 23227). Sample concentrations and volumes were normalised to load 300 µg protein onto each sucrose gradient, with 10 % of the sample set aside as inputs. Ultracentrifugation was performed for 16 h at 158000 g (30400 rpm) at 4°C using an SW-41-Ti rotor on a Sorvall WX+ Ultra Series instrument. Following ultracentrifugation, samples were fractionated into 1.5 mL Eppendorf tubes using a BioComp Gradient Fractionator. Fractions were subsequently concentrated using Triochloroacetic (TCA) precipitation. 100 % TCA (Sigma, 92128-100G) was added to each fraction (10 % of the fraction volume added), the samples gently inverted to mix and samples incubated on ice for 15 min. Samples were then centrifuged at high speed (17000 g, 10 min, 4°C), the supernatant removed and samples incubated with 500 µL ice-cold 100 % acetone (Roth, 9372.1) for 10 min on ice. Samples were once more centrifuged at high speed, the supernatant removed and washed once more with ice-cold acetone. After centrifugation and removal of the supernatant, the resultant pellet was resuspended in 2X Laemmli buffer (containing 20 % Tris HCl pH 8.0 (Thermo Fisher, AM9855G)) and boiled for 10 min at 95 °C, before analysis of the fractions by western blotting.

### Western blotting

Protein extracts were mixed with Laemmli buffer, boiled for 10 minutes at 95 °C, and centrifuged for 1 min at 13000 g to remove aggregates. Samples were resolved on a 4-12% gradient Bis-Tris gel (Bio-Rad, 3450125), and transferred to a PVDF membrane using the Trans Blot Turbo system (Bio Rad). Blots were incubated in blocking buffer (5% milk in PBS-T (0.05 % Tween 20 (Sigma, P9416-100ML)) for > 30 min at room temperature with shaking. Primary antibodies were diluted in blocking buffer and incubated for > 16 h at 4°C with shaking. Following primary antibody incubation, samples were then washed 3 x 15 min with PBS-T at room temperature with shaking, before incubation with an anti-rabbit HRP-conjugated secondary antibody (Cell Signaling, 7074S) at 1:1000 dilution for 1 h at room temperature. Samples were then washed 3 x 15 min with PBS-T. Blots were developed by incubation with Clarity Max™ ECL Substrate (Bio-Rad, 1705062) and then visualised on a Bio-Rad Chemidoc imager. Antibodies used for Western blotting and their working dilutions can be found in **Key Resources Table.**

### TMT-expression proteomics

#### Sample preparation - FASP + C18 cleanup

Cells were lysed in lysis buffer (2% SDS, 50 mM HEPES, 1 mM PMSF, 1 x protease inhibitor cocktail from Roche). Samples were resuspended and incubated at room temperature for 20 minutes before they were sonicated in a Branson ultrasonic processor with microtip on ice (0.5 seconds on / 0.5 seconds off, 30 seconds total, 20% input). The lysate was centrifuged at 16,000 g for 10 minutes at 20°C and the supernatants transferred to a fresh tube. The protein concentration was measured with a Pierce BCA Protein Assay (Product Nr.: 23227) according to the manufacturer instructions. The FASP protocol was employed for proteolytic digestion of as previously reported (Wiśniewski *et al*, 2009) with slight modifications. Briefly, DTT (83.3 mM) was added to the lysis buffer and the sample incubated at 95°C for 5 minutes. Microcon 30 Ultracel YM-30 filter units were primed once with 200 µL urea buffer (8 M urea in 100 mM Tris/HCl, pH 8.5). Samples (100 µg) were then loaded onto the filter units at 14,000 rcf at 20°C for 15 minutes, washed with UA solution (8 M Urea in 100 mM Tris/HCl, pH 8.5). Alkylation was performed with 200 µl 50 mM Iodacetamide in UA solution. Samples were again centrifuged (14,000 rcf, 10 minutes) after a 30-minute incubation period in the dark. Next, samples were washed twice with 100 µl UA and then twice with 100 µl 50 mM TEAB. Samples were then digested in 40 µl 50 mM TEAB, pH 8.5, at a protein to enzyme ratio of 50:1. The spin columns were sealed and incubated at 37 °C overnight. On the next day, samples were spun down (14,000 rcf, 20 minutes) and the columns were washed once with 40 µL 50 mM TEAB and once with 50 µL 0.5M NaCl. The combined filtrate was acidified with 30% TFA until a pH below 3 was achieved. Samples were then subjected to C18 cleanup with Pierce™ Peptide Desalting spin columns according to the manufacturer’s instructions (ThermoFisher Scientific; Catalog number 89851, 89852).

#### Peptide quantification

Desalted peptides were reconstituted in LC-grade water, and peptide concentrations were determined using the Pierce™ Quantitative Fluorometric Peptide Assay (Thermo Fisher Scientific, Ref: 23290) according to manufacturer’s instructions. Fluorescence measurements were performed on a SpectraMax® i3x Multi-Mode Microplate Reader (Molecular Devices).

#### TMT labelling

Peptides were subsequently buffered to a final concentration of 50 mM HEPES, pH 8.5. Labelling was performed at a peptide concentration of ≥ 1 µg/µL using TMT reagent prepared in anhydrous acetonitrile at 16.7 µg/µL, applied at a TMT-to-peptide mass ratio of approximately 2:1 to 3:1. The final acetonitrile concentration during the labelling reaction was 35% (v/v). Samples were incubated for 1 hour at room temperature with gentle agitation (∼600 rpm). Labelling reactions were subsequently quenched by the addition of 5% hydroxylamine solution in HEPES buffer to achieve a final hydroxylamine concentration of 0.2–0.4%, followed by incubation for 15 minutes at room temperature. Labelled peptides were pooled, dried by vacuum centrifugation, and reconstituted in 0.1% trifluoroacetic acid for subsequent processing. Labelling efficiency was routinely assessed following final analysis.

#### Offline fractionation of TMT pool

Pooled TMT labelled sample was concentrated and desalted by using a Pierce Peptide Desalting Spin Columns (Catalog Number 89851 and 89852) according to manufacturer’s instructions with minor modifications: Peptide elution was done in 2 steps: 40% ACN, 100mM TEAB and 70% ACN, 100 mM TEAB solution. Eluates were pooled and brought to dryness in a speedvac vacuum concentrator. Dry peptides were resuspended in 30ul 10mM ammonium formate (pH 10) and fractionated by reverse-phase chromatography at basic pH by using a Gemini-NX C18 (150C×C2 mm, 3 μm, 110 A,) column (Phenomenex, Torrance, USA) on an Ultimate 3000 RSLC micro system (Thermo Fisher Scientific) equipped with a fraction collector. Peptides were separated at a flow rate of 50 µl/min in 10 mM ammonia formate buffer (pH 10.0) and eluted over a 70 min nonlinear gradient from 0 to 100% solvent B (90% acetonitrile, 10 mM ammonium formate, pH 10.0). Thirty-six concatenated fractions were collected in a time-based manner (at 30 s intervals) between minute 11.5 and 57. Fractions were immediately acidified by adding 5 µl 30% TFA. After fractionation organics were removed in a vacuum concentrator and sample were stored at -20C until MS analysis.

#### LC-ESI-MS/MS data acquisition

For MS analysis fractions were reconstituted in 150 µl 0.1% TFA and 5 µl was injected onto the machine. For proteomic data acquisition, a nanoflow LC−ESI-MS/MS setup comprised of a Dionex Ultimate 3000 RSLCnano system coupled to a Fusion Lumos mass spectrometer (both ThermoFisher Scientific Inc.) was used in positive ionisation mode. MS data acquisition was performed in data-dependent acquisition (DDA) mode. For proteome analyses, peptides were delivered to a trap column (Acclaim™ PepMap™ 100 C18, 3 μm, 5 × 0.3 mm, Thermo Fisher Scientific) at a flowrate of 5 μL/min in HPLC grade water with 0.1% (v/v) TFA. After 10 min of loading, peptides were transferred to an analytical column (ReproSil Pur C18-AQ, 3 μm, Dr. Maisch, 500 mm × 75 μm, self-packed) and separated using a stepped gradient from minute 11 at 8% solvent B (0.4% (v/v) FA in 90% ACN) to minute 61 at 28% solvent B and minute 81 at 40% solvent B at 300 nL/min flow rate. The nano-LC solvent A was 0.4% (v/v) FA HPLC-grade water. MS1 spectra were recorded at a resolution of 60,000 using an automatic gain control target value of 4 × 10^5 and a maximum injection time of 50 ms. The cycle time was set to 2 seconds. Only precursors with charge state 2 to 6 which fall in a mass range between 360 to 1300 Da were selected and dynamic exclusion of 30 s was enabled. Peptide fragmentation was performed using higher energy collision dissociation (HCD) and a normalised collision energy of 35%. The precursor isolation window width was set to 1.3 m/z. MS2 spectra were acquired at a resolution of 30,000 with an automatic gain control target value of 5 × 10^4 and a maximum injection time of 54 ms. Using synchronous precursor selection, the top 10 fragment ions of the MS2 scans were isolated and subjected to HCD fragmentation in the linear ion trap using 55% normalized collision energy. MS3 spectra were acquired in the Orbitrap at a resolution of 50,000 over a scan range of 100 to 1000 Th, using an AGC target of 1e5 and a maximum injection time of 120 ms. The reagent tag type in the filer IsobaricTagExclusion was set to TMTpro.

#### Data analysis

For all DDA measurements, MaxQuant (version 2.6.7.0) with its built-in search engine Andromeda (Cox *et al*, 2011; Tyanova *et al*, 2016a) was used for peptide identification and quantification. MS2 spectra were searched against all Swiss-Prot canonical protein sequences obtained from UniProt (UP000005640, downloaded: 19 March 2025), supplemented with common contaminants (built-in option in MaxQuant). Trypsin/P was specified as the proteolytic enzyme. Precursor tolerance was set to 4.5 ppm, and fragment ion tolerance to 20 ppm. The minimal peptide length was defined as seven amino acids, and the “match-between-run” function was disabled. Quantification was performed on the MS3 level. “18plex” (TMTpro) was selected as isobaric labels and the reporter mass tolerance was set to 0.003 Da. For proteome analyses, carbamidomethylated cysteine was set as a fixed modification and oxidation of methionine and N-terminal protein acetylation as variable modifications. The FDR was set to 100%. This search results were then used as input files for Oktoberfest (v.0.8.3) (Picciani *et al*, 2024). We performed Prosit rescoring (Gessulat *et al*, 2019) and quantification via the picked-group-FDR approach (v.0.8.1) (The *et al*, 2022). The Prosit models Prosit_2020_irt_TMT for retention time prediction and Prosit_2020_intensity_TMT for intensity prediction were employed. The output files were filtered at 1% FDR. Perseus was used for data analysis (Tyanova *et al*, 2016b). Briefly, “common contaminants”, “reversed”, and “only identified by site” were filtered out, the intensities for each TMT channel were log2-transformed, and median-centric normalised. Samples were then categorically annotated to group replicates together and the “Hawaii plot” function with default values was performed. The obtained matrix was exported for further analyses.

### Materials availability

All reagents generated in this study are available upon request to the lead contact with a completed materials transfer agreement.

### Data and code availability

CUT&RUN datasets generated in this study can be accessed on the Gene Expression Omnibus (GEO) database under the series accession number GSE300600. Raw microscopy data will be provided by the corresponding authors on request. The IPython notebooks used to perform analysis of data generated within this manuscript will be made available upon publication (https://github.com/seruggialab). All mass spectrometric raw data files as well as the data analysis output files have been deposited to the ProteomeXchange Consortium (http://proteomecentral.proteomexchange.org) (Vizcaíno *et al*, 2014) via the PRIDE partner repository with the dataset identifier PXD065443.

## Supporting information

Key Resources Table

## ACKNOWLEDGMENTS

We thank the CCRI FACS Core Unit for assistance with flow cytometry and cell sorting, Miriam Abele and the Proteomics team of the Molecular Discovery Platform at CeMM for mass-spectrometric data acquisition, the CeMM Biomedical Sequencing Facility, and the Core Imaging Facility at the Medical University of Vienna for technical support. Finally, we thank members of the Seruggia lab for providing important feedback on this project, and Stefan Kubicek, David Haselbach and Zuzana Hodáková for insightful discussions. This research was funded in whole or in part by the Austrian Science Fund (FWF) [10.55776/P36069]. For open access purposes, the author has applied a CC BY public copyright license to any author accepted manuscript version arising from this submission. Research in the Seruggia laboratory is supported by the Austrian Science Fund (FWF, projects P36069 and P36302), the European Research Council (ERC) under the Horizon 2020 research and innovation programme (grant agreement 947803) and under Marie Skłodowska–Curie (grant agreement 101061151). G.S.-F. is supported by the Austrian Academy of Sciences. Aithyra and the Winter lab are supported by the Boehringer Ingelheim Foundation and the Austrian Academy of Sciences. The Winter lab is further supported by funding from the European Research Council (ERC) under the European Union’s Horizon 2020 research and innovation programme (grant agreement 851478), by team KOODAC via the Cancer Grand Challenges partnership funded by Cancer Research UK (CGCATF-2023/100013), Institut National Du Cancer (INCa) and KiKa (Children Cancer Free Foundation), as well as by funding from the Austrian Science Fund (FWF, projects P5918723, P36746 and P7909) and the Vienna Science and Technology Fund (WWTF, project LS21-015).

## AUTHOR CONTRIBUTIONS

Conceptualization, P.B. and D.S.; investigation, P.B., D.S.; formal analysis, P.B., H.B., C.S., A.P.K.; methodology, P.B., H.B., C.S., G.O., M.Z., S.M.; resources, G.E.W., D.S.; validation, P.B. H.B., S.M.; writing – original draft, P.B and D.S.; writing – review and editing, P.B., H.B., C.S., G.E.W., G.S.-F. and D.S.; funding acquisition, D.S.; supervision, G.E.W., G.S.-F. and D.S.

## Supplemental information

Figures S1–S4 and Tables S1

Table S2. Excel file containing additional data too large to fit in a PDF, related to Figure 5.

Table S3. Excel file containing additional data too large to fit in a PDF, related to Figures 1, 2, 4, 5, S1, S2, S4.

## SUPPLEMENTARY FIGURES

**Figure S1.**
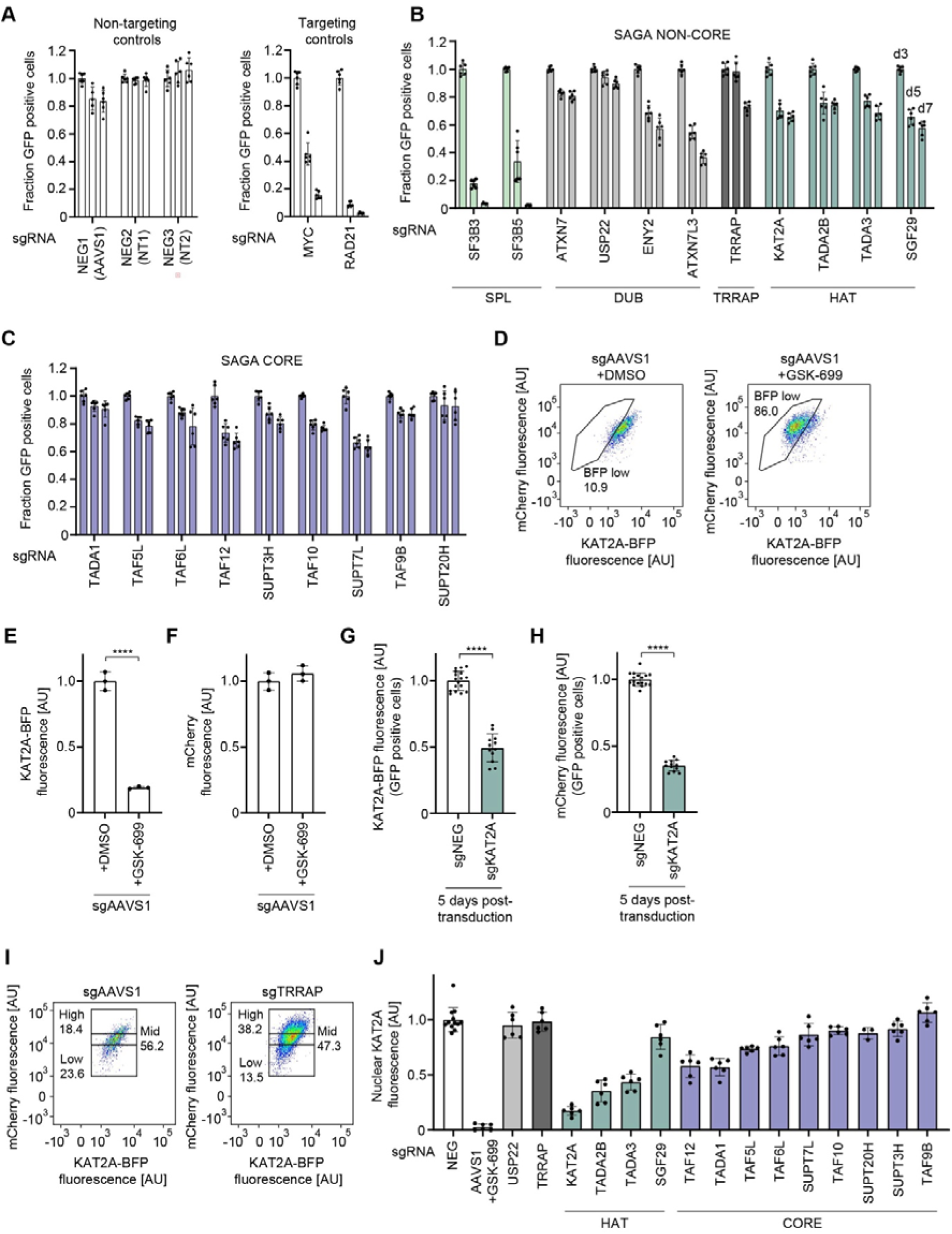
Arrayed CRISPR screening to monitor KAT2A stability upon knockout of SAGA components; related to Fig. 1. **A-C)** To assess gRNA dropout of our arrayed SAGA-focused library, the fraction of GFP positive cells for each guide was measured 3-, 5-, and 7-days post-transduction of wild type HAP1^Cas9^ cells expressing the KAT2A stability reporter. The fraction of GFP positive cells on each day was normalised to the day 3 value for each guide. One gRNA per SAGA subunit is plotted. Dots represent the fraction of GFP positive cells for technical replicates on each day, error bars indicate the standard deviation, bars indicate the mean for each condition. **A)** Fraction of GFP positive cells for targeting and non-targeting control gRNAs. **B)** Fraction of GFP positive cells for non-CORE module SAGA subunits. **C)** Fraction of GFP positive cells for CORE module SAGA subunits. **D-F)** Validation of the responsiveness of the KAT2A stability reporter to chemical perturbation. Wild type HAP1^Cas9^ cells expressing the KAT2A stability reporter were transduced with GFP expressing gRNAs targeting AAVS1. 5 days post-transduction cells were treated with DMSO or 100 nM GSK-699 for 6 h, before analysis by flow cytometry. Dots in bar plots represent the mean normalised KAT2A-BFP or mCherry fluorescence of technical replicates, error bars indicate the standard deviation, bars indicate the mean for each condition. Significance was tested using a two-tailed unpaired t-test. **D)** 2D flow cytometry plots of KAT2A-BFP vs mCherry fluorescence for wild type HAP1^Cas9^ cells expressing the KAT2A stability reporter and transduced with gRNAs against AAVS1, after treatment with DMSO or GSK-699, as indicated. **E)** Mean KAT2A-BFP fluorescence of AAVS1 transduced cells treated with DMSO or GSK-699, analysed by flow cytometry. **F)** Mean mCherry fluorescence of AAVS1 transduced cells treated with DMSO or GSK-699, analysed by flow cytometry. **G, H)** Validation of the responsiveness of the KAT2A stability reporter to genetic perturbation of KAT2A. Wild type HAP1^Cas9^ cells expressing the KAT2A stability reporter were transduced with GFP expressing gRNAs targeting AAVS1 or KAT2A. 5 days post-transduction cells were analysed by flow cytometry. Dots represent the mean normalised KAT2A-BFP or mCherry fluorescence of technical replicates, error bars indicate the standard deviation, bars indicate the mean for each condition. Significance was tested using a two-tailed Mann–Whitney U-test. **G)** Mean KAT2A-BFP fluorescence of AAVS1 or KAT2A transduced cells analysed by flow cytometry. **H)** Mean mCherry fluorescence of AAVS1 or KAT2A transduced cells analysed by flow cytometry. **I)** 2D flow cytometry plots of KAT2A-BFP vs mCherry fluorescence for wild type HAP1^Cas9^ cells expressing the KAT2A stability reporter, 5 days after transduction with gRNAs against AAVS1 or TRRAP, as indicated. Black boxes mark mCherry low, mid, and high populations for each condition, as indicated. **J)** Quantification of nuclear KAT2A fluorescence for experimental conditions as indicated, as in **Fig. 1E**. Dots represent the median nuclear KAT2A fluorescence of technical replicates, error bars indicate the standard deviation, bars indicate the mean for each condition. Data information: **D)** (****) P < 0.0001); two-tailed unpaired t-test. **F, G** (****) P < 0.0001; two-tailed Mann–Whitney U-test. Biologically independent replicates: **(A-C, G-J)** (n = 2). Technical replicates: **D-F** (n = 3). In **D-I**, only GFP positive cells were analysed. In **E, F,** the data was normalised relative to the mean KAT2A-BFP or mCherry fluorescence of AAVS1 transduced cells treated with DMSO. In **G, H,** the data was normalised relative to the mean KAT2A-BFP or mCherry fluorescence of cells transduced with negative control guides. The number of cells analysed by microscopy in **J** is listed in **Table S3.**

**Figure S2.**
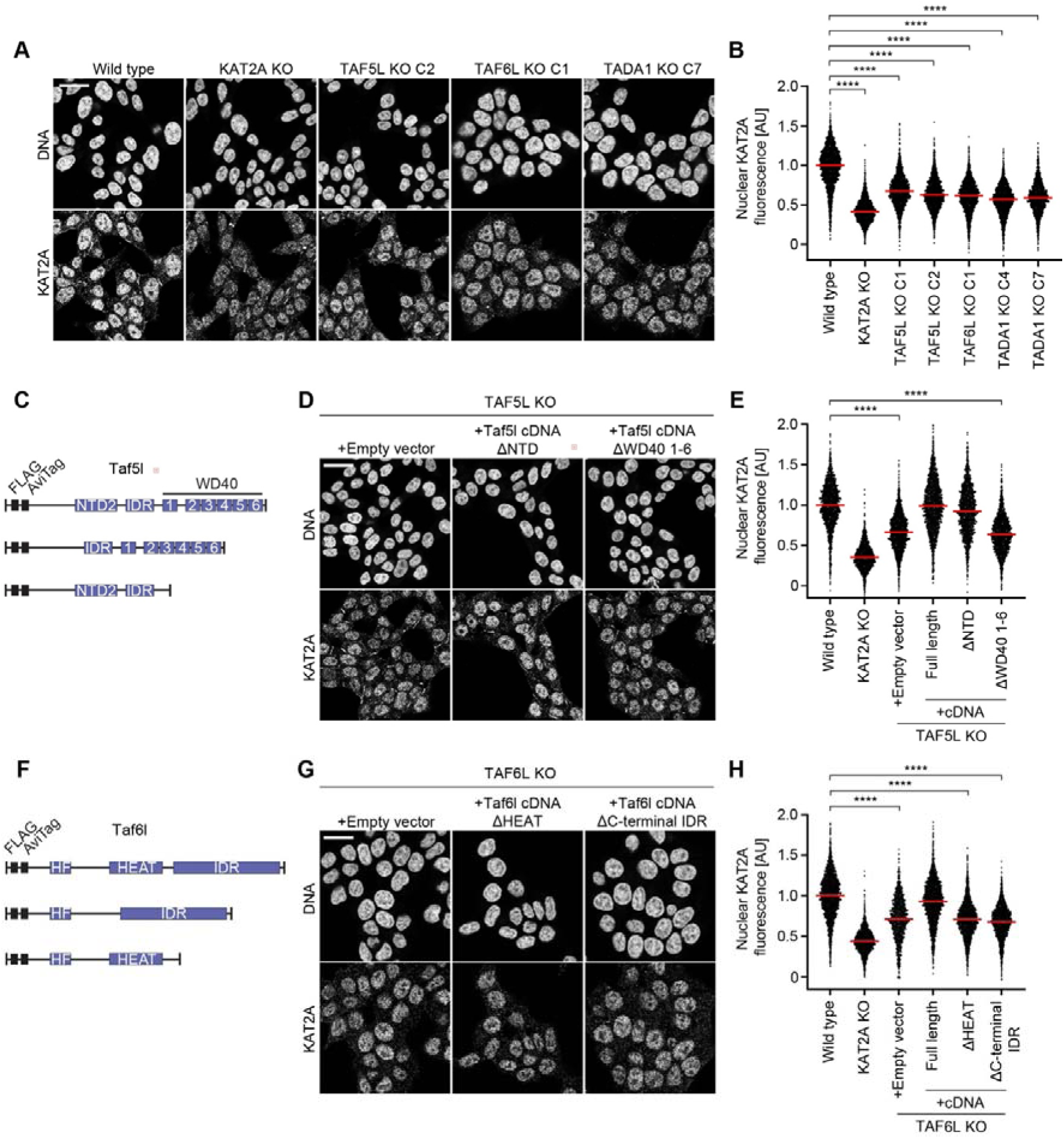
Components of the SAGA CORE regulate KAT2A protein abundance and HAT function; related to Fig. 2. **A-B)** Knockout of TAF5L, TAF6L, or TADA1 reduces nuclear KAT2A levels. **A)** Representative immunofluorescence images of wild type, KAT2A KO, TAF5L KO, TAF6L KO, or TADA1 KO cells shown as indicated. Cells were fixed for immunofluorescence and KAT2A was stained with an anti-KAT2A antibody. DNA was stained with Hoechst 33342. **B)** Quantification of nuclear KAT2A fluorescence by immunofluorescence for wild type and CORE KO HAP1 cells as indicated. Dots represent the mean nuclear KAT2A fluorescence of individual cells; red bars indicate the mean. Two independent clones shown for TAF5L KO and TADA1 KO cells. Significance was tested using a two-tailed Mann–Whitney U-test. **C)** Schematic of TAF5L cDNA overexpression constructs used in **Fig 2A, B, S2D, E.** cDNA for full-length murine TAF5L, or TAF5L lacking its NTD2 domain or C-terminal WD-40 domains were overexpressed in TAF5L KO cells. **D)** Representative immunofluorescence images of TAF5L KO overexpressing murine cDNA overexpression constructs as indicated. Cells were fixed for immunofluorescence and KAT2A was stained with an anti-KAT2A antibody. DNA was stained with Hoechst 33342. **E)** Quantification of nuclear KAT2A fluorescence by immunofluorescence for conditions as indicated. Dots represent the mean nuclear KAT2A fluorescence of individual cells; red bars indicate the mean. Significance was tested using a two-tailed Mann–Whitney U-test. **F)** Schematic of TAF6L cDNA overexpression constructs used in **Fig 2A, B, S2G, H.** cDNA for full-length murine TAF6L, or TAF6L lacking its HEAT domain or a C-terminal intrinsically disordered region (IDR) was overexpressed in TAF6L KO cells. **G)** Representative immunofluorescence images of TAF6L KO overexpressing murine cDNA overexpression constructs as indicated. Cells were fixed for immunofluorescence and KAT2A was stained with an anti-KAT2A antibody. DNA was stained with Hoechst 33342. **H)** Quantification of nuclear KAT2A fluorescence by immunofluorescence for conditions as indicated. Dots represent the mean nuclear KAT2A fluorescence of individual cells; red bars indicate the mean. Significance was tested using a two-tailed Mann–Whitney U-test. Data information: **B, E, H)** (****) P < 0.0001; two-tailed Mann–Whitney U-test. Biologically independent replicates: **(A, B, D, E)** (n = 3), **G, H** (all conditions n = 3 apart from TAF6L KO^empty^ ^vector^ (n = 2)). In **B** the data was normalised relative to the median nuclear KAT2A fluorescence of wild type HAP1 cells. In **E, H,** the data was normalised relative to the mean nuclear KAT2A fluorescence of wild type HAP1 cells. All images show single Z-slices. Scale bars: 20 µm. The number of cells analysed by microscopy in **B, E, H,** is listed in **Table S3.**

**Figure S3.**
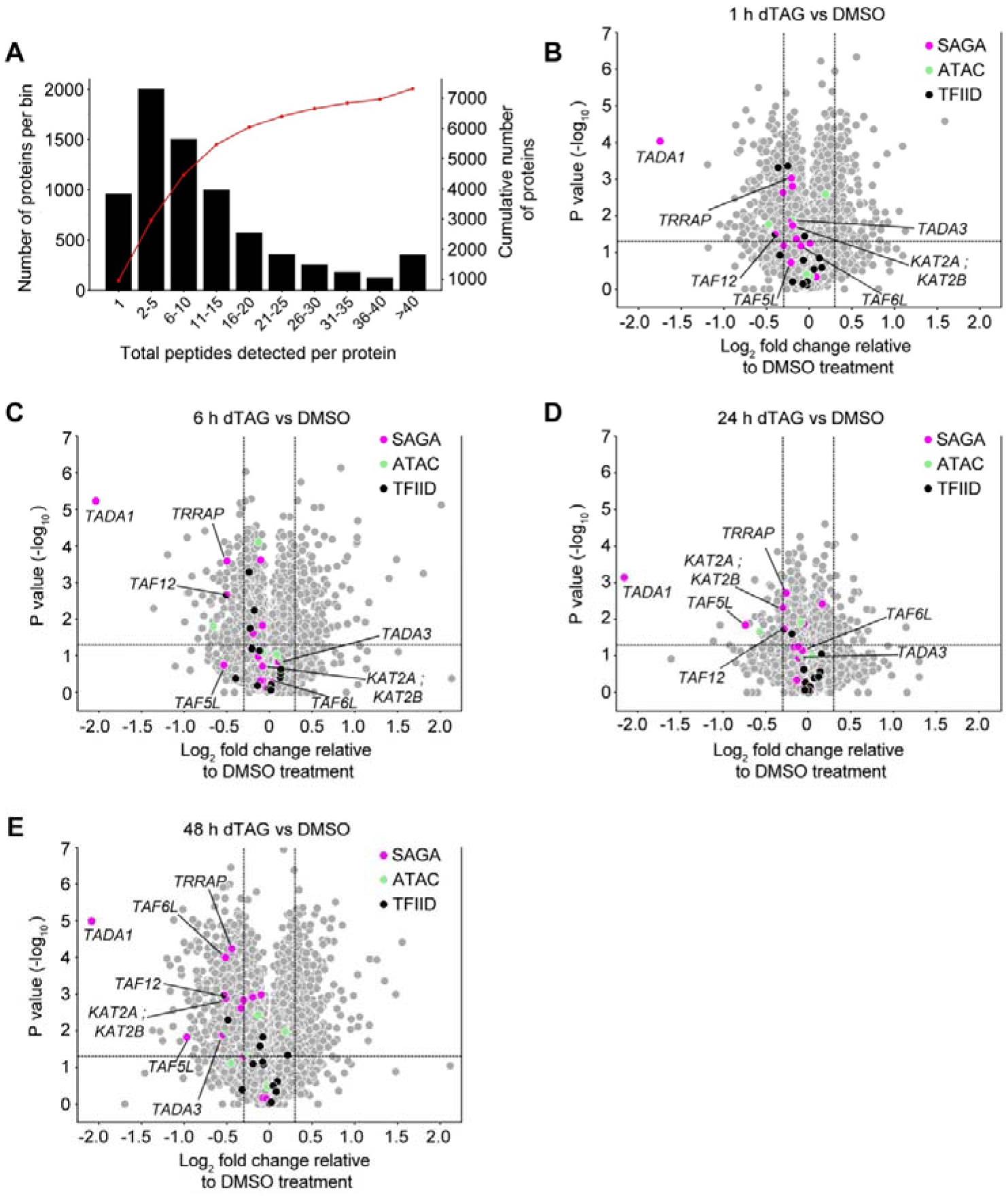
Progressive destabilisation of SAGA CORE and HAT upon depletion of TADA1; related to Fig. 4. **A)** Binned peptide counts for each protein detected in the TMT-expression dataset, plotted as a histogram. The number of proteins within each bin is plotted on the left Y-axis. The cumulative number of identified proteins is plotted on the right Y-axis (red line). **B – E)** Volcano plots of differentially abundant proteins following acute depletion of TADA1 for the indicated time points in TADA1-dTAG HAP1 cells. Fold changes and p-values were calculated by comparison of dTAG^v^-1 treated cells for each time point with DMSO-treated controls. Significantly downregulated proteins: Log_2_ fold change ≤ 0.3 and -log_10_ p-values ≥ 1.3. Magenta dots represent SAGA complex components. Green dots represent ATAC complex components. Black dots represent TFIID complex components. Subunits common to more than one complex are labelled with both colours. Components of the SAGA complex with significantly reduced abundance after 24 h and/or 48 h dTAG treatment are labelled with black lines. Data information: Biologically independent replicates: **A-E** (n = 3).

**Figure S4.**
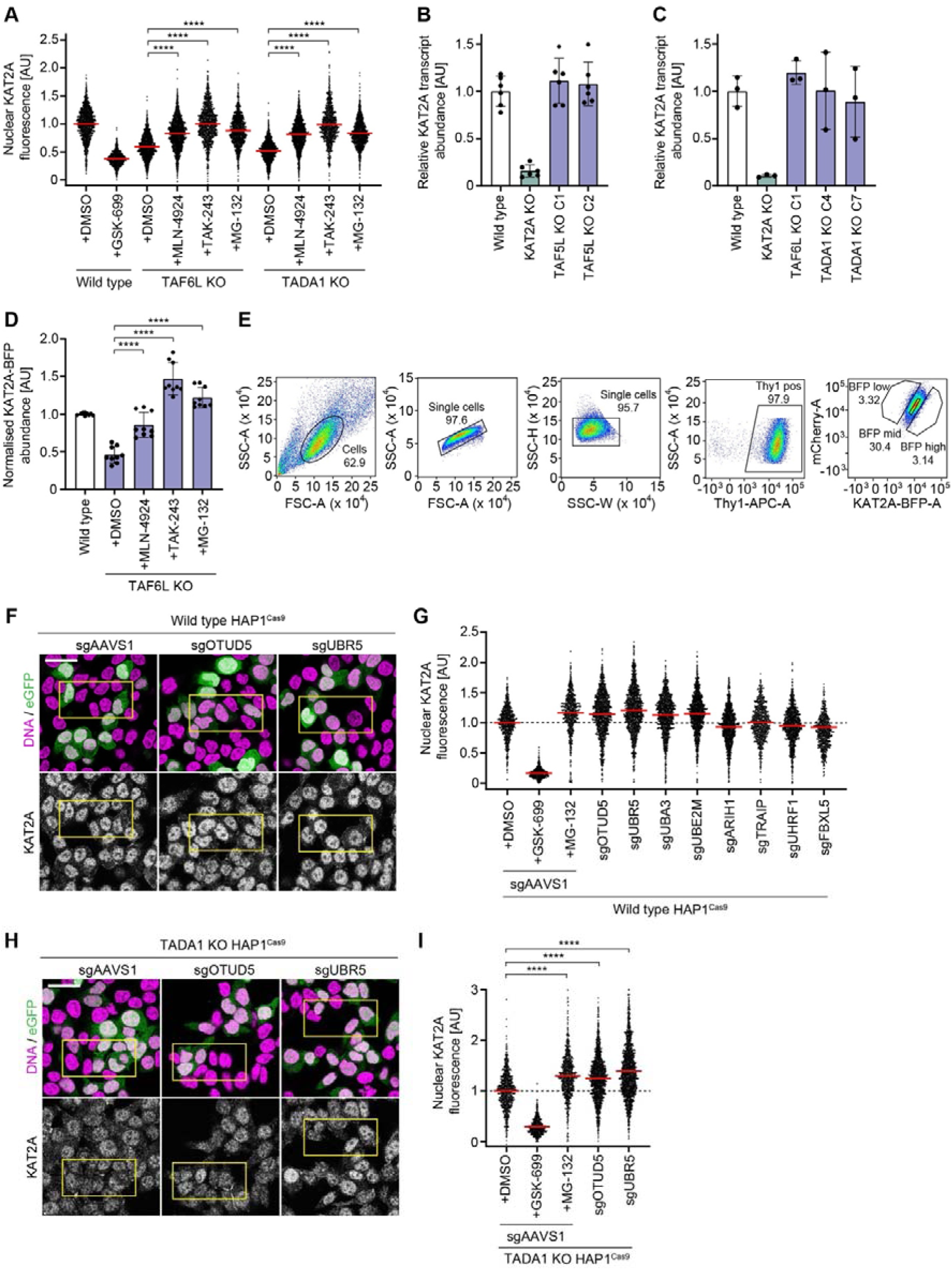
Non-complexed KAT2A is degraded in a process mediated by UBR5 and OTUD5; related to Fig. 5. **A)** Quantification of nuclear KAT2A fluorescence by immunofluorescence following treatment with inhibitors as indicated. Wild type, TAF6L KO, and TADA1 KO cells were treated with DMSO, GSK-699, MLN-4924, TAK-243, or MG-132 as in Fig. 5A, B before fixation for immunofluorescence. Following compound treatment, cells were fixed and KAT2A was stained with an anti-KAT2A antibody. Dots represent the mean nuclear KAT2A fluorescence of individual cells; red bars indicate the mean. Significance was tested using a two-tailed Mann–Whitney U-test. Quantification for wild type HAP1 cells treated with DMSO and GSK-699 is the same as in Fig. 5B to allow side-by-side comparison with the knockout cells treated with inhibitors. **B)** RT**-**qPCR of KAT2A mRNA transcript levels in wild type, KAT2A KO, and TAF5L KO (two independent clones) HAP1 cells. mRNA levels were normalised to GAPDH. **C)** RT**-**qPCR of KAT2A mRNA transcript levels as in **B** for wild type, KAT2A KO, TAF6L KO, and TADA1 KO (two independent clones) HAP1 cells. **D)** Quantification of normalised KAT2A-BFP fluorescence using the KAT2A stability reporter. Wild type and TAF6L KO cells expressing the KAT2A stability reporter were treated with inhibitors as in Fig. 5C before analysis by flow cytometry. Dots represent the mean normalised KAT2A-BFP fluorescence of technical replicates, error bars represent the standard deviation, bars indicate the mean for each condition. Significance was tested using a two-tailed Mann–Whitney U-test. **E)** Representative plots of the gating strategy used for FACS in the pooled CRISPR screen presented in Fig. 5E. Numbers indicate the percentage of cells inside the respective gate. TAF5L KO HAP1^Cas9^ cells expressing the KAT2A stability reporter were gated using the following logic: 1) FSC-A vs SSC-A to gate cells, 2) FSC-A vs FSC-H and SSC-H vs SSC-W to gate single cells, FSC-A vs Thy1-APC-A to identify sgRNA expressing cells, and then KAT2A-BFP-A vs mCherry-A to sort KAT2A-BFP^low^, KAT2A-BFP^mid^, and KAT2A-BFP^high^ populations. The gating for KAT2A-BFP was adjusted dynamically to sort the top and bottom 3 % of KAT2A-BFP expressing cells. **F-I)** Immunofluorescence validation of UPS effectors whose knockout increases KAT2A abundance in TADA1 KO^Cas9^ cells and wild type^Cas9^ HAP1 cells, as in Fig. 5F, G. 5 days post-transduction, cells were fixed and KAT2A was stained with an anti-KAT2A antibody. DNA was stained with Hoechst 33342. Where indicated, cells were treated with GSK-699 (100 nM) or MG-132 (1 µM) for 8 h before fixation. All other samples were treated with DMSO. **F)** Representative immunofluorescence images of wild type HAP1^Cas9^ cells transduced with gRNAs against AAVS1, OTUD5, or UBR5, as indicated. **G)** Quantification of nuclear KAT2A fluorescence of wild type HAP1^Cas9^ cells transduced with the indicated gRNAs. GFP positive cells were analysed. The data for two independent gRNAs is merged for each UPS gene. Dots represent the mean nuclear KAT2A fluorescence of individual cells; red bars indicate the mean. Significance was tested using a two-tailed Mann–Whitney U-test. **H)** Representative immunofluorescence images of TADA1 KO HAP1^Cas9^ cells transduced with gRNAs against AAVS1, OTUD5, or UBR5, as indicated. **I)** Quantification of nuclear KAT2A fluorescence of TADA1 KO HAP1^Cas9^ cells transduced with the indicated gRNAs. GFP positive cells were analysed. The data for two independent gRNAs is merged for each UPS gene. Dots represent the mean nuclear KAT2A fluorescence of individual cells; red bars indicate the mean. Significance was tested using a two-tailed Mann–Whitney U-test. Data information: **C, D, G, I** (****) P < 0.0001; two-tailed Mann–Whitney U-test. Biologically independent replicates: **A** (n = 6), **B-D** (n = 3), **F-I** (n = 2). In **A, B,** the data was normalised relative to the mean KAT2A transcript abundance of wild type HAP1 cells. In **C** the data was normalised relative to the mean nuclear KAT2A fluorescence of wild type HAP1 cells treated with DMSO. In **G, I** only GFP positive cells were analysed, and data was normalised relative to the mean nuclear KAT2A fluorescence of AAVS1 transduced cells treated with GSK-699 for the respective cell line. All images show single Z-slices. Yellow boxes indicate inset regions showing non-transduced (GFP negative) and transduced (GFP positive) cells. Scale bars: 20 µm. The number of cells analysed by microscopy in **G, I** is listed in **Table S3.**

